# Identification of Two Elusive Human Ribonuclease MRP-Specific Protein Components

**DOI:** 10.1101/2025.01.19.633795

**Authors:** Rui Che, Bhoomi Mirani, Monireh Panah, Xiaotong Chen, Hong Luo, Andrei Alexandrov

## Abstract

All known protein components of one of the longest-studied human ribonucleoprotein ribozyme nuclear Ribonuclease MRP (RNase MRP), which processes pre-rRNA at ITS1 site 2, are shared with Ribonuclease P (RNase P), which cleaves pre-tRNA 5′ leader sequences. Our genome-wide forward genetic screening identified two poorly characterized human genes, which we named RPP24 and RPP64. We show that these two genes are required for pre-rRNA ITS1 site 2 processing and their protein products efficiently associate with RNA MRP. Unlike all other human RNase MRP protein components, RPP24 and RPP64 are not required for RNase P activity and do not associate with RNase P-specific RNA H1. Despite extremely limited sequence homology, RPP24 and RPP64 exhibit predicted structural similarities to two RNase MRP-specific components in *S. cerevisiae*, with specific differences in RPP64 regions of substrate recognition. Collectively, our functional screening and validation revealed the first two protein components unique to human nuclear RNase MRP.

## INTRODUCTION

Human Ribonuclease P (RNase P) and nuclear Ribonuclease MRP (RNase MRP) are some of the longest-studied ribonucleoprotein ribozymes. They are composed of protein subunits and a distinct catalytic non-coding RNA, whose discovery in RNase P earned Sydney Altman the 1989 Nobel Prize, which he shared with Thomas Cech^1,2^. Whereas RNase P is primarily responsible for processing the 5′-leader sequences of pre-tRNAs^1,3,4^ and tRNA-like structures^5-7^ and is conserved across all three domains of life^8^, RNase MRP is found only in eukaryotes^9,10^ and is known for the initial processing of precursor rRNA (pre-rRNA) at a specific site, namely ITS1 site 2 in humans^11^ and ITS1 site A3 in yeast *S. cerevisiae*^12,13^. Besides processing pre-rRNA, RNase MRP is known to cleave additional RNA substrates, including generating primers for mitochondrial DNA replication^14,15^ and processing the 5′ UTR of cyclin B2^16^ and CTS1 mRNA^17^. Mutations in the RNA moiety of human RNase MRP are associated with genetic disorders such as cartilage-hair hypoplasia (CHH) and anauxetic dysplasia, both of which are classified as ribosomopathies, i.e., resulting from defects in ribosome biogenesis^18,19^. Whereas the structure of the human RNase MRP complex is unknown, recent structures of yeast RNase P^20^ and RNase MRP^21,22^ as well as human RNase P^23^ (**Fig. 1A, B,** and **D**) revealed their organization and catalytic mechanisms^9^, as well as critical substrate recognition determinants, including the key roles of yeast RNase MRP-specific protein subunits in recognition of the pre-rRNA substrate^21^.

**Figure 1.**
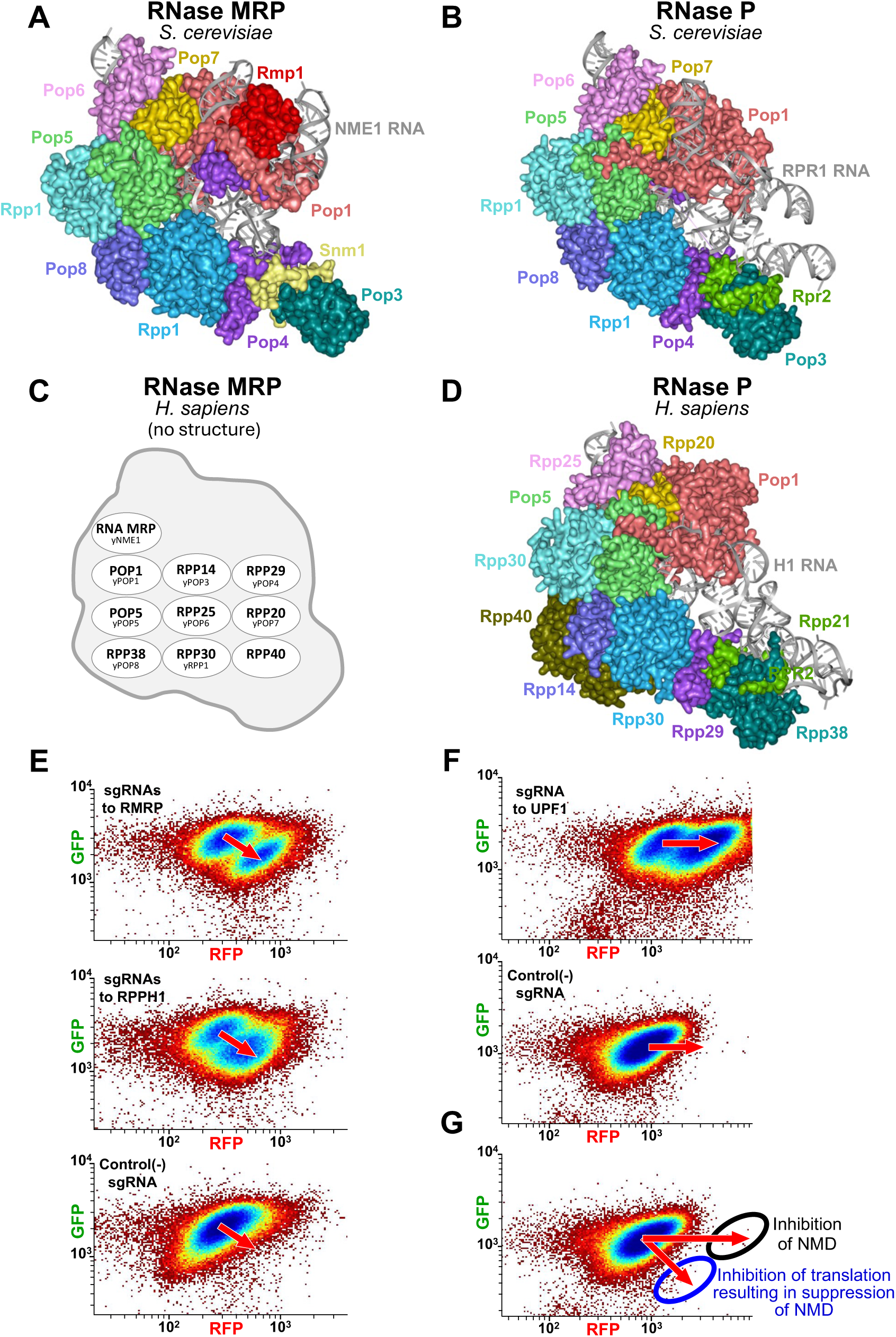
RNase MRP/P complexes and use of the Fireworks fluorescence system to detect their deficiency in single human cells. RNase MRP and RNase P share most of their protein components. Human RNase MRP has no known unique protein components. **A-D.** Known components and structures of *S. cerevisiae* and human RNase MRP and RNase P. **A.** PDB: 7C79^21^. **B.** PDB: 6AGB^20^. **C.** The structure of human RNase MRP remains unknown^9^. **D.** PDB: 6AHR^23^. **E.** Knockouts of the human RNase MRP/P-specific RNA genes, RMRP and RPPH1, induce right-and-down fluorescence shift in the Fireworks cell line **F.** Knockout of the NMD factor UPF1 induces a rightward fluorescence shift in the Fireworks cell line, as reported previously^30^. **G.** Schematic of the different responses of the Fireworks cells to (i) NMD inhibition and (ii) translation inhibition, which suppresses NMD due to NMD dependence on active translation^31-35^. (Also see **Fig. S1**).

In yeast, there are two such RNase MRP-specific protein subunits, Snm1 and Rmp1, which are absent in RNase P. The first subunit, Snm1, assists in the folding of the yeast RNase MRP RNA, NME1, to form the substrate-binding module^9,21,22^. The second subunit, Rmp1, together with Pop4 and Pop1, coordinates the single-stranded pre-rRNA substrate and its characteristically flipped cytosine base, enabling RNase MRP’s substrate specificity^21,22^. The additional eight protein subunits of yeast RNase MRP—Pop1, Pop3, Pop4, Pop5, Pop6, Pop7, Pop8, and Rpp1—are all shared with RNase P^9,21,22,24^. Furthermore, the trimer Pop1-Pop6-Pop7 is also found in the yeast telomerase holoenzyme^25^, where it is required for the correct localization of the telomerase RNA, TLC1^26^.

In contrast, in humans, since the discovery of the human RNase MRP over 35 years ago^27,28^, no RNase MRP-specific protein subunits have been identified^11,24^. It has therefore remained a mystery whether such subunits exist^9,11^, as the nine proteins known to associate with human RNase MRP— Pop1, Pop5, Rpp14, Rpp20, Rpp25, Rpp29(Pop4), Rpp30, Rpp38, and Rpp40—are also known to be components of RNase P^24,29^ (**Fig. 1C, D**). The lack of known RNase MRP-specific protein components has greatly hindered studies of human RNase MRP, including the identification of its substrates and substrate specificity determinants, understanding its function, and determining its structure, which remains unknown^9,11^.

Using a genome-wide forward genetic screening that leveraged the requirement of RNase MRP and RNase P for reporter translation, we identified two poorly characterized yet highly enriched human genes, c18orf21 and c3orf17, which we named RPP24 and RPP64, respectively. We demonstrate that these two genes are specifically required for the activity of human RNase MRP but not RNase P. The protein products of RPP24 and RPP64 associate extensively and exclusively with the only known human RNase MRP-specific component, RNA MRP, but not with the RNase P-specific RNA H1. Our findings establish RPP24 and RPP64 as the first two human protein components unique to RNase MRP.

## RESULTS

### Forward genetic screening for factors impacting human translation enriched components of Ribonuclease MRP and Ribonuclease P, as well as ribosome biogenesis and translation initiation factors

To identify factors affecting the essential human ribonuclear complexes nuclear RNase MRP and RNase P, which are indispensable for translation due to their established roles in processing the pre-rRNA ITS1 site 2^11^ and pre-tRNA 5′-leader sequences^1,3,11^, we performed a genome-wide CRISPR sgRNA-based forward genetic screening that detects translation defects using a dual-fluorescence reporter system. This system can distinguish, in high throughput, defects in translation from defects in transcription, which are otherwise indistinguishable using a basic fluorescent protein reporter, as for such a reporter both defects produce a decrease in fluorescence.

As shown previously^30^ and illustrated for UPF1 in **Fig. 1F**, the Fireworks cells (**Fig. S1A**) exhibit a rightward fluorescence shift upon the knockout of *bona fide* NMD components (**Fig. 1F, G**). In contrast, as observed previously^30^, translation defects, including those resulting from knockouts of lncRNA MRP and lncRNA H1 (the catalytic components of RNase MRP and RNase P, respectively), induce a right-and-downward fluorescence shift (**Fig. 1E, G** and **S1C**). This right-and-downward shift occurs because the inhibition of translation reduces the susceptibility of the premature termination codon (PTC)-containing RFP reporter (**Fig. S1A**) to NMD, as NMD requires active protein synthesis^31-35^. Consequently, for the same-cell PTC-containing reporter, translation inhibition and the resulting suppression of NMD lead simultaneously to (i) reduced translation of the reporter and (ii) an increase in its PTC-containing mRNA available for translation. Together, these two opposing effects of translation inhibition on the PTC-containing RFP Fireworks reporter result in an overall increase in its fluorescence, whereas the fluorescence of the PTC-lacking GFP Fireworks reporter decreases (**Figs. 1G** and **S1A**, **B**). This response, previously used to distinguish human *bona fide* NMD factors^30^, enables efficient FACS-based isolation of cells with NMD-impacting defects in translation, differentiating translation defects from those in both transcription and NMD (**Fig. S1B**).

To minimize undesired enrichment of guide RNAs targeting NMD factors in the forward genetic screen that detects translation defects *via* their impact on NMD, we constructed a genome-wide lentiviral “omission” CRISPR library consisting of 90,260 guide RNAs targeting 20,861 human genes and deliberately lacking guide RNAs targeting known *bona fide* NMD factors (**Fig. 2A** and **Supplemental Data 1**).

**Figure 2.**
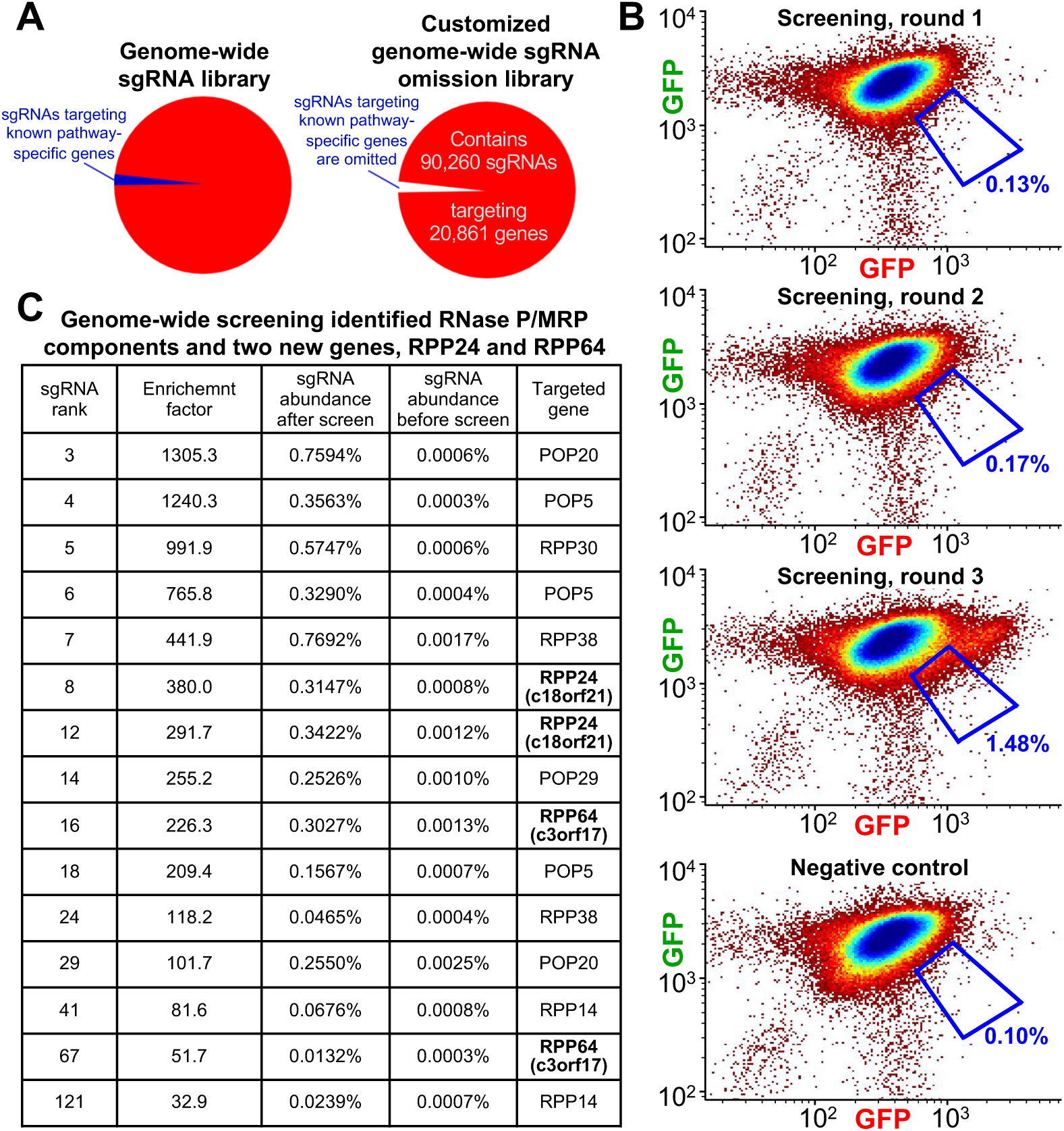
Genome-wide iterative forward genetic screening identified components of RNase MRP and RNase P, as well as two poorly characterized genes. **A.** Schematic of the customized lentiviral genome-wide CRISPR-Cas9 sgRNA omission library used in the screening. **B.** FACS-sorting rounds of iterative genome-wide screening enrich Fireworks cells in the right-and-down blue sorting gate. **C.** Enrichment of two poorly characterized genes c3orf17, c18orf21, and shared components of RNase MRP/P. (Also see **Fig. S2** and **Supplemental Data S1**.)

We conducted three rounds of genome-wide iterative FACS-based forward genetic screening (**Figs. 2B** and **S2**) of the NMD omission library (**Figs. 2A**). During these screening rounds, the Fireworks cells harboring guide RNAs that cause a right-and-downward fluorescence shift (**Fig. 1G**) were enriched in the blue sorting gate (**Fig. 2B**). The enrichment levels were 0.13%, 0.17%, and 1.48% (**Fig. 2B**) after the first, second, and third rounds, respectively (**Fig. S2**).

Deep sequencing of guide RNAs isolated from cells enriched after the third screening round (**Fig. 2B**) revealed that guide RNAs targeting components shared by RNase P and RNase MRP, including POP20, POP5, RPP30, RPP38, POP29, and RPP14, display some of the highest enrichment (**Fig. 2C** and **Supplemental Data 1**). This high enrichment can be attributed to the concurrent disruption of two pathways indispensable for translation—tRNA processing and pre-rRNA processing—the combined effect of which would be expected to produce a major two-pronged impact on translation and, consequently, NMD (**Fig. 2C** and **Supplemental Data 1**).

Consistent with the enrichment of factors impacting translation, the screening additionally enriched guide RNAs targeting tRNA synthetases, including the highly enriched YARS (YARS1) and QARS (QARS1); a component of the eukaryotic translation initiation factor 3 (eIF-3) complex, EIF3I; components of the small subunit (SSU) processome, such as PWP2 (UTP1), IMP3, IMP4, and RPS19BP1 (AROS); ribosome biogenesis factors NOC4L (UTP19), NOL12, and NOM1; ribosomal proteins RPS28, RPS16, and RPS11; and numerous other ribosome biogenesis factors (**Supplemental Data 1**).

Strikingly, guide RNAs targeting two poorly characterized human genes, c18orf21 and c3orf17, exhibited enrichment as high as that of components shared by both RNase MRP and RNase P (**Fig. 2C** and **Supplemental Data 1**).

### RPP24 and RPP64 are specifically required for the activity of human nuclear RNase MRP, and not RNase P

Individual FACS-based validation of guide RNAs targeting the screening-enriched c18orf21 (RPP24) and c3orf17 (RPP64) as well as nuclear RNase MRP and RNase P components RMRP RNA, POP5, RPP14, RPP20, RPP29, RPP30, RPP38, RPP40, RPP21, and RPPH1 confirmed the right-and-downward fluorescence shift (**Fig. 3A** and **S3B**) in the Fireworks cell line (**Fig. S1A, B**), indicating a translation-mediated impact of their knockouts on NMD (**Fig. 1G**). Due to these similarities, and the strikingly high enrichment of c18orf21 (RPP24) and c3orf17 (RPP64), comparable to that of the components of nuclear RNase MRP and RNase P (**Fig. 2C** and **Supplemental Data 1**), we assessed the effects of the knockouts of c18orf21 (RPP24) and c3orf17 (RPP64), obtained as shown in **Fig. S3A**, on the activities of human nuclear RNase MRP and RNase P. Unlike viability-based approaches, fluorescence-based enrichment of knockout cells enables their efficient isolation well in advance of the onset of lethality, facilitating the analysis of early knockout effects. As measured using RT-qPCR across the ITS1 site 2^11^ of the endogenous rRNA precursor, knockouts of c18orf21 (RPP24) and c3orf17 (RPP64), quantified in **Figs. 3D** and **S3B**, produced respective 5.9- and 11.6-fold increase in the levels of the unprocessed pre-rRNA precursor (**Fig. 3B**, left graph). This increase was consistent with the 6.9-, 4.6-, 6.8-, 6.9-, 7.0-, 5.0-, 6.9, and 4.5-fold increase in the levels of the ITS1 site 2-unprocessed pre-rRNA precursor resulting from the knockouts of RMRP RNA, POP5, RPP14, RPP20, RPP29, RPP30, RPP38, and RPP40, respectively (**Fig. 3B**, left graph; the knockouts are quantified in **Fig. 3D** and **S3B, C**). To rule out off-target effects of RPP24- and RPP64-targeting sgRNAs on processing of the ITS1 site 2 of the endogenous pre-rRNA precursor, we confirmed that additional sgRNAs toward RPP24 and RPP64 replicated this effect (**Figs. 3B**, right graph, and **S3B**). As negative controls, knockouts of RPP21 and RPPH1, the unique protein and RNA subunits of the human RNase P, respectively, produced no effects on the cleavage of the endogenous pre-rRNA ITS1 site 2 (**Fig. 3B**).

**Figure 3.**
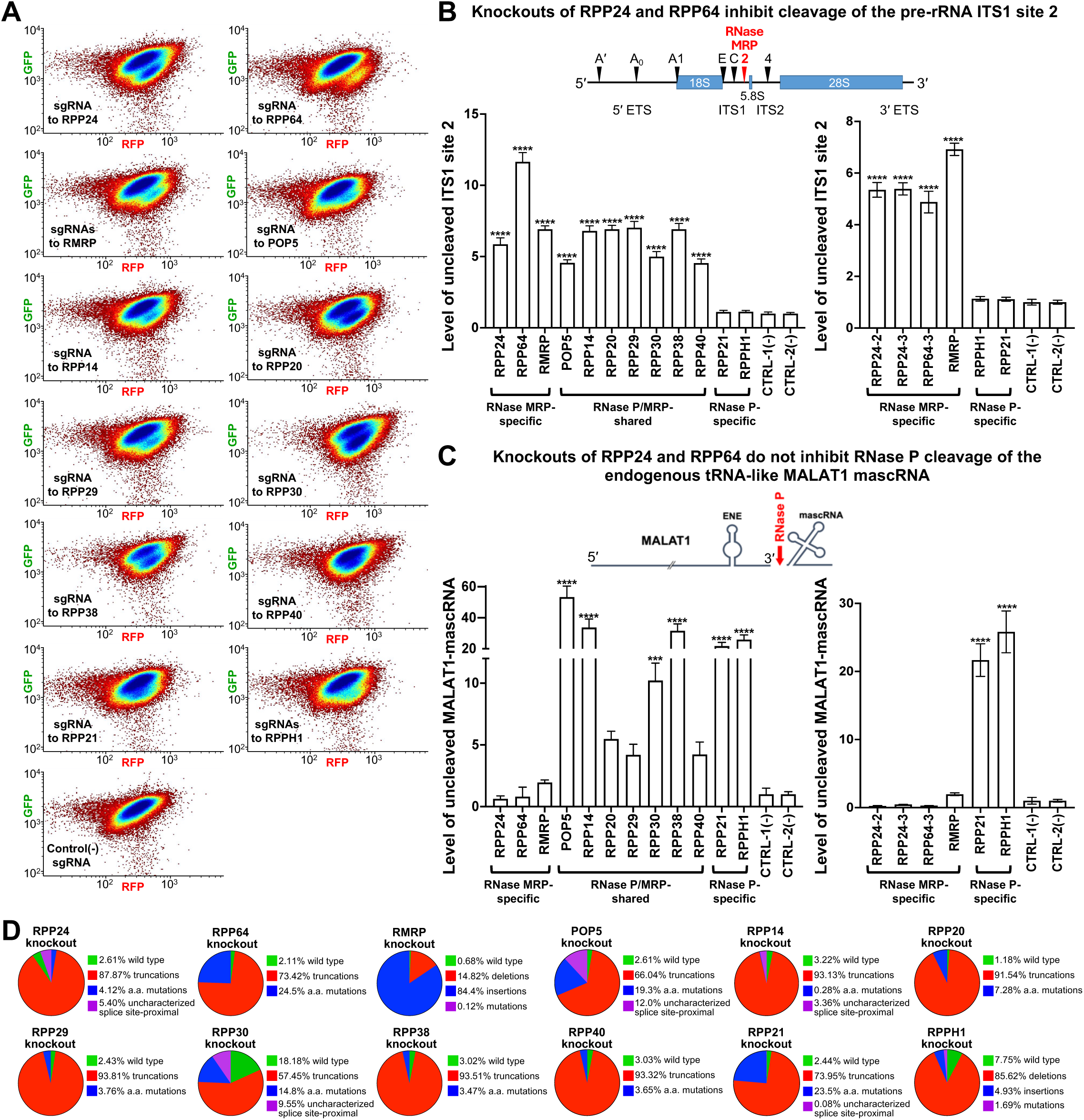
Human RPP24 and RPP64 are required for the cleavage of endogenous substrates of RNase MRP but not RNase P. **A.** Knockouts of RPP24, RPP64, and components of RNase MRP/P produce right-and-down shift in fluorescence of the Fireworks cells. **B.** Knockouts of RPP24, RPP64, and RNase MRP components result in pre-rRNA processing defect at ITS1 site 2. **C.** Unlike knockouts of RNase P-specific and RNase P/MRP-shared components, knockouts of RPP24 and RPP64 do not cause a defect in the cleavage of tRNA-like mascRNA of the endogenous lncRNA MALAT1. RT-qPCRs are described in Methods Details; data are presented as means of at least n=3 replicates; error bars represent standard deviation. **D.** Analysis of CRISPR knockouts (obtained as shown in **Fig. S3A**) used in panels A-C, using Illumina sequencing of the sgRNA-targeted genomic sites. (Also see **Fig. S3** and **Supplemental Data S4**)

In contrast to the impact of the knockouts of RPP24 and RPP64 on the processing of the pre-rRNA ITS1 site 2, the same knockouts produced no increase in the levels of the endogenous RNase P-unprocessed MALAT1-mascRNA precursor (**Fig. 3C**, left graph). Consistently, additional sgRNAs toward RPP24 and RPP64 confirmed the lack of an increase (**Fig. 3C**, right graph). For comparison, the knockouts of the RNase P/MRP-shared components POP5, RPP14, RPP20, RPP29, RPP30, RPP38, and RPP40, as well as the knockouts of the RNase P-specific components RPP21 and RPPH1 resulted in a 4.2- to 53-fold increase in the levels of the RNase P-unprocessed endogenous lncRNA MALAT1 (**Fig. 3C**). These findings confirm the efficiency of the knockouts (quantified in **Fig. 3D**, and **S3B, C**) and are consistent with the essential roles of POP5, RPP14, RPP20, RPP29, RPP30, RPP38, RPP40, RPP21, and RPPH1^9,23^ in the activity of human endogenous RNase P.

Collectively, our data demonstrate that RPP24 and RPP64 are specifically required for the activity of human nuclear RNase MRP in pre-rRNA processing but are dispensable for the cleavage of the endogenous tRNA-like MALAT1 mascRNA by RNase P.

### Human Rpp24 and Rpp64 specifically co-purify with the only known RNase MRP-specific component, RNA MRP, but not with the RNase P-specific RNA component, RNA H1

Since both RPP24 and RPP64 are specifically required for the activity of human RNase MRP and not RNase P (**Fig. 3B, C**), we asked whether Rpp24 and Rpp64 associate with RNA MRP in human cells. We immunopurified these proteins from nuclear extracts and quantified the amounts of RNA MRP and RNA H1 in the eluted fractions.

As shown in the top panel of **Figure 4A**, FLAG purification of Rpp24 and Rpp64 resulted in 131- and 246-fold enrichment of RNA MRP, respectively, compared to that of the “No ORF” and Mettl1-FLAG^36^ negative controls. The enrichment of RNA MRP in the Rpp24 and Rpp64 fractions exceeded that observed with the RNase P-specific protein component Rpp21 (46-fold) and was comparable to the enrichment seen with the RNase P/MRP-shared components, Rpp25 (191-fold).

**Figure 4.**
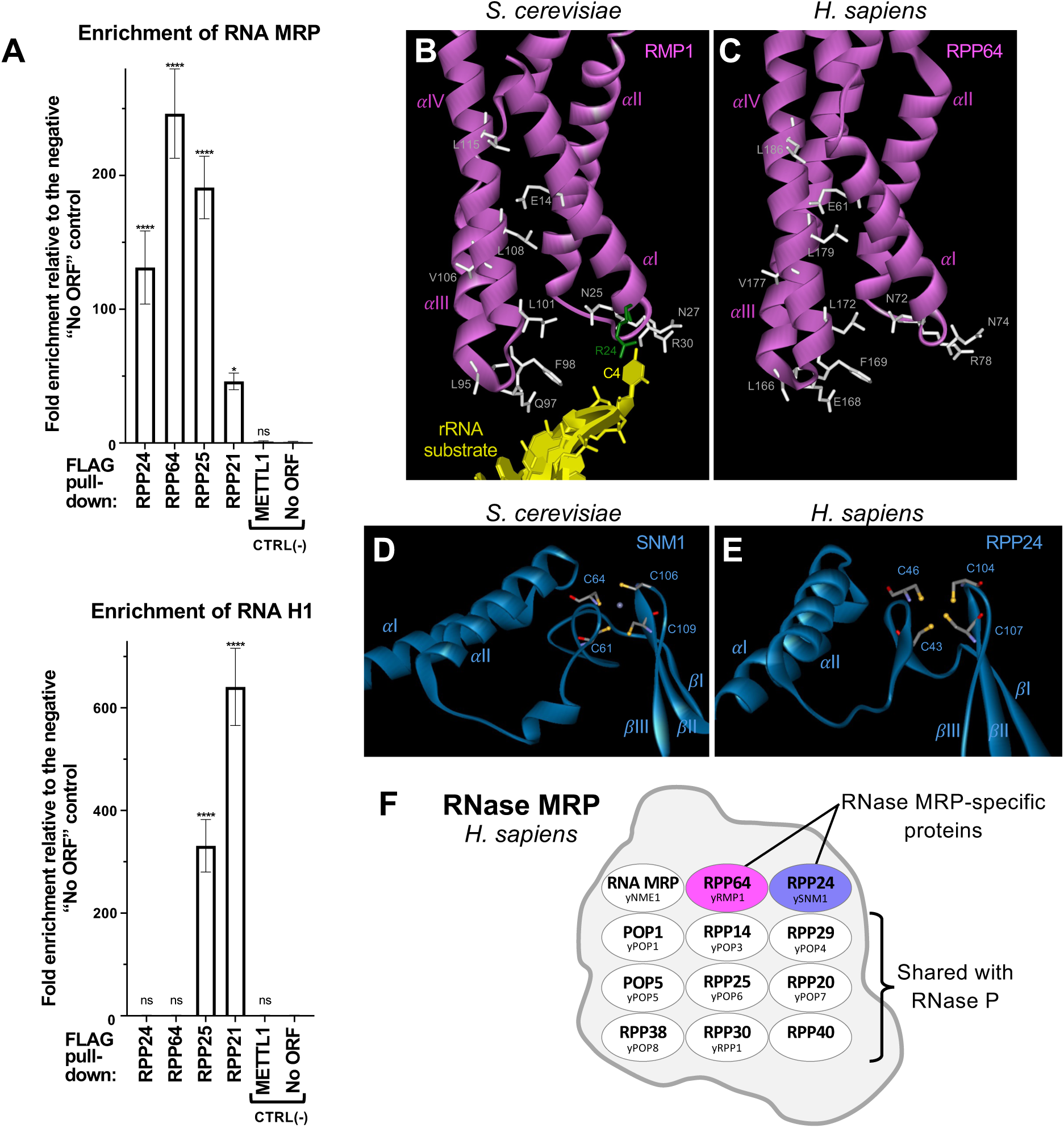
Human Rpp24 and Rpp64 specifically associate with RNA MRP and exhibit structural similarity to yeast RNase MRP-specific components. **A.** FLAG-tagged human Rpp24 and Rpp64 specifically co-purify the only known RNase MRP-specific component, RNA MRP, but not the RNase P-specific RNA component, RNA H1 from HEK293T cells. Fold enrichment of RNA MRP relative to “No ORF” negative control in the FLAG-purified fractions of Rpp24, Rpp64, Rpp25, Rpp21, Mettl1, and “No ORF control” are 131.2±27.4, 246.3±33.4, 191.0±23.4, 46.08±6.20, 0.999±0.62, and 1.00±0.21, respectively. Fold enrichment of RNA H1 relative to “No ORF” negative control in the FLAG-purified fractions of Rpp24, Rpp64, Rpp25, Rpp21, Mettl1, and “No ORF control” are 0.94±0.21, 0.97±0.21, 331.1±51.3, 640.9±75.3, 0.86±0.18, and 1.00±0.19, respectively. (Shown as means±SD.) Mettl1-FLAG^36^ represents a non-interacting negative control. **B-E.** Human Rpp64 and Rpp24 exhibit predicted structural similarities to the key pre-rRNA substrate recognition region of Rmp1, and the C2C2 zinc finger region of Snm1, respectively, in the cryo-EM structure of *S. cerevisiae* RNase MRP. **B.** PDB: 7C7A^21^. **C.** AlphaFold DB: AF-Q6NW34-F1-v4^54^. **D.** PDB: 7C7A^21^. **E.** AlphaFold DB: AF-Q32NC0-F1-v4^54^. **F.** Rpp24 and Rpp64 represent the first human nuclear RNase MRP-specific protein components that are absent in RNase P. (Also see **Fig. S4**.)

By contrast, the same immunopurifications of Rpp24 and Rpp64 (**Fig. 4A**, compare bottom and top panels) did not enrich RNA H1 compared to that of the “No ORF” negative control. In comparison, immunopurification of the RNase P-specific protein component Rpp21 and the RNase P/MRP-shared component Rpp25 resulted in 640- and 331-fold enrichment of RNA H1, respectively, reflecting their known association with RNA H1 in the RNase P complex.

Together, these findings demonstrate a specific and preferential enrichment of Rpp24 and Rpp64 with RNA MRP, the only known RNase MRP-specific component, and no enrichment with RNA H1, the RNase P-specific RNA component. These results suggest that Rpp24 and Rpp64 represent previously unrecognized protein components of human RNase MRP that are not shared with RNase P.

### Human c3orf17 (RPP64) displays limited sequence homology but shares predicted structural similarities with *S. cerevisiae’s* RNase MRP-specific component Rmp1 in its key pre-rRNA substrate recognition region

Whereas Protein BLAST^37^ for human Rpp64 yielded negligible alignment score (E-value of 0.61) with *S. cerevisiae*’s RNase MRP-specific component Rmp1 (**Fig. S4A**), the cryo-EM structure of *S. cerevisiae*’s Rmp1 and the AlphaFold-predicted structure of Rpp64 displayed the following similarities (**Fig. 4B, C**): First, the overall arrangement of the four N-terminal alpha-helices (αI – αIV) comprising amino acids 5-123 in yeast Rmp1 and 52-200 in human RPP64 is similar (**Fig. 4**, compare panels **B** and **C**). Second, twelve amino acids conserved between human Rpp64 and yeast Rmp1 (**Fig. S4A**) exhibit similar positions and orientations (**Fig. 4B, C**; shown in white). Third, several of these conserved amino acids—Asn72 (Asn25), Asn74 (Asn27), Arg78 (Arg30), Leu166 (Leu95), Glu168 (Gln97), Phe169 (Phe98), and Leu172 (Leu101)—are located in the loops connecting alpha-helices αI and αII (Loop I-II) and αIII and αIV (Loop III-IV) (**Fig. 4B, C**). Critically, in the structures of yeast RNase MRP^21,22^, these Rmp1 loops I-II and III-IV represent key elements of substrate recognition: together with residues of Pop1 and Pop4, they coordinate the ITS1 substrate backbone phosphates of C_2_ and A_3_^21^, enabling the flipped state of the substrate nucleotide C_4_ (**Fig. 4B**). Nevertheless, unlike yeast Rmp1, in which Arg24 and Gln28 in loop I-II and Gln97 in loop III-IV are critical for coordinating the flipped state of the substrate nucleotide C_4_ in the yeast RNase MRP structure^21^ (**Fig. 4B**), human Rpp64 carries Ser71, Arg75, and Glu168 at the corresponding positions, potentially reflecting species-specific requirements for recognizing and cleaving distinct pre-rRNA ITS1 sequences^11,38^.

### Human c18orf21 (RPP24) displays limited sequence homology but shares predicted structural similarities with *S. cerevisiae* RNase MRP-specific components Snm1

Although Protein BLAST^37^ for human Rpp24 produced negligible alignment score (E-value of 0.76) with *S. cerevisiae*’s RNase MRP-specific component Snm1 (**Fig. S4B**), the cryo-EM structure of *S. cerevisiae*’s Snm1 and the AlphaFold-predicted structure of Rpp24 displayed notable structural similarities: First, the N-terminal regions of Snm1 and Rpp24 have an overall αα-loop-βββ arrangement (**Fig. 4**; compare panels **D** and **E**). Specifically, they are comprised of two α-helices, αI and αII, spanning amino acids 5-46 in yeast Snm1 and 3-33 in human Rpp24, followed by a single α-helical turn-containing loop formed by amino acids 47-92 in Snm1 and 34-54 in Rpp24, and three β-sheets (βI – βIII) comprised of amino acids 93-117 in yeast Snm1 and 55-114 in human Rpp24 (**Fig. 4D, E**). Second, both human and yeast proteins contain four conserved cysteines arranged in a CXXC…CXXC motif (**Fig. S4B**). In the structure of yeast Snm1, these four cysteines coordinate Zn^+2^, forming an experimentally determined C2C2-type zinc finger^21,39^ (**Fig. 4D**). AlphaFold predicts a similar zinc-finger arrangement for Rpp24 (**Fig. 4E**). In both, the cryo-EM structure of yeast Snm1 and the AlphaFold-predicted structure of Rpp24, one cysteine C106 (C104) is located within βII, another (C109 in Snm1; C107 in Rpp24) is located in the βII – βIII turn, and the remaining two cysteines (C61 and C64 in Snm1; C43 and C46 in Rpp24) reside in the αII – βI loop.

Consistent with our functional and pull-down analyses, the structural similarities of human Rpp24 and Rpp64 to the only two known yeast RNase MRP-specific (i.e., not present in RNase P) protein components, Snm1 and Rmp1 (**Fig. 4B-E**), further support the RNase MRP-specific roles of human Rpp24 and Rpp64.

## DISCUSSION

To uncover additional human genes essential for the roles of RNase MRP and RNase P in rRNA biogenesis and tRNA processing, we leveraged the requirement of both rRNA biogenesis and tRNA processing for translation. We conducted forward genetic screening to identify genes whose knockouts disrupt reporter translation to the major degree observed with knockouts of components of RNase MRP and RNase P. To enable high-throughput screening for translation defects, we employed a dual-fluorescence reporter system^30^ that uses translation-mediated inhibition of NMD to rapidly distinguish defects in translation from defects in transcription^31-34^ (**Figs. S1B** and **1E-G**). To minimize unintended enrichment of *bona fide* NMD factors, we employed a genome-wide lentiviral omission library that deliberately lacked guide RNAs targeting known NMD components. Our genome-wide iterative screening identified two poorly characterized genes, c18orf21 and c3orf17 (we named RPP24 and RPP64, respectively), that exhibited some of the highest enrichment (**Fig. 2C**), typical of the components shared by nuclear RNase MRP and RNase P (**Fig. 2C**).

Until our identification of RPP24 and RPP64, it remained a puzzle whether any human factors specific to human RNase MRP exist^9,11,24^. Our characterization of RPP24 and RPP64 revealed that: First, CRISPR sgRNA-based knockouts of RPP24 and RPP64 result in accumulation of the ITS1 site 2-unprocessed pre-ribosomal RNA (**Fig. 3B**). In contrast, the same knockouts of RPP24 and RPP64 have no effect on RNase P cleavage of the tRNA-like mascRNA of the endogenous lncRNA MALAT1 (**Fig. 3C**), suggesting their RNase MRP specificity. Second, pull-downs of Rpp24 and Rpp64 specifically enrich the catalytic RNA component of RNase MRP, RNA MRP (**Fig. 4A**, top panel), but not the RNase P catalytic RNA component, RNA H1 (**Fig. 4A**, bottom panel), suggesting RNase MRP-specific association. Third, despite very low sequence homology (**Fig. S4**), both human Rpp24 and Rpp64 exhibit predicted structural similarities to the only two known *S. cerevisiae*’s RNase MRP-specific components Snm1 and Rmp1, respectively (**Fig. 4B-E**). Remarkably, the structural (**Fig. 4B, C**) and sequence (**Fig. S4A**) similarities of human Rpp64 are most pronounced in the region of yeast Rmp1 that enables the flipped state of the pre-rRNA substrate nucleotide C_4_ (**Fig. 4B**), which is one of the most distinctive features of the yeast RNase MRP’s substrate recognition and specificity^9,21^.

Supporting our finding of RPP64’s role in RNase MRP-mediated pre-rRNA ITS1 site 2 processing, Rpp64(c3orf17/NEPRO^40^) is known to localize in the nucleolus^41,42^, and its mice knockout exhibits mis-localization of 18S rRNA^41^. Mass spectrometry identified RNase P/MRP-shared subunits in Rpp64 preparations^42^ and, separately, Rpp24, Rpp64, and RNase P/MRP components in preparations of MOB3C^43^. RPP64 is linked to cartilage hair hypoplasia (CHH) and anauxetic dysplasia^44,45^, rare disorders, which are also associated with mutations in RMRP^18^ (RNase MRP-specific RNA) and POP1^46^ (a shared RNase P/MRP protein). Genome-wide co-essentiality mapping and supervised machine learning predicted the association of RPP24(c18orf21) and RPP64 with RNase P/MRP complexes^47,48^. Separately, recent genome-scale computational analysis of perturbative maps of transcriptional and morphological data also suggested RPP24’s involvement in the RNase MRP complex^49^.

In addition to establishing the roles of RPP24 and RPP64, our findings confirm that human RPP29(POP4), RPP14, POP5, RPP20, RPP30, RPP38, RPP40, and RNA MRP are required for pre-rRNA ITS1 site 2 cleavage (**Fig. 3B**). We cannot draw such conclusions for POP1 due to insufficient knockout efficiency (data not shown) and for RPP25, which shows no effect (data not shown), possibly due to its redundancy with RPP25L, as reported for RNase P^50,51^. Consistent with reports of Rpp21’s interaction with human RNA MRP *in vitro*^52,53^, Rpp21-FLAG co-purifies a significant amount of RNA MRP—46 times that of the negative controls (**Fig. 4A**, top panel). Nevertheless, consistent with human RPP21 being specifically required for activity of RNase P, we confirm that RPP21 is not required for the ITS1 site 2 cleavage (**Fig. 3B**).

In summary, we demonstrate the existence of two human nuclear MRP-specific protein components (**Fig. 4F**) that have eluded identification for over three decades. Their discovery greatly simplifies the study of RNase MRP independently from RNase P, which was previously hindered by the shared nature of all RNase MRP protein subunits. Our identification of RPP24 and RPP64 paves the way for studying substrate specificity determinants of RNase MRP, uncovering its additional potential endogenous human RNA substrates, further delineating its cellular function, and contributing to its future structural studies.

## Supporting information

Supplemental Data Descriptions

## DATA AVAILABILITY

The high-throughput sequencing data generated in this study, including CRISPR sgRNA library screening datasets have been deposited to NCBI Sequence Read Archive (SRA), accession number: PRJNA1204971, and will be released by the time of publication.

## ACKNOWLEDGMENTS

We thank Kevin Babson and Greenwood Genetic Center for Illumina NextSeq sequencing support and Krista Knowles for help with library construction. This work was supported by National Institutes of Health grant GM139769 (project 1) to Andrei Alexandrov.

## AUTHOR CONTRIBUTIONS

R.C. and A.A. conceived and performed experiments and wrote the manuscript, B.M. and M.P. performed experiments, X.C. and H.L. provided bioinformatic data analyses, AA supervised the project.

## METHODS DETAILS

### Construction of plasmids

#### LentiCRISPR plasmids expressing individual guide RNAs

Guide RNA-containing 140-base-pair PCR products were amplified using primers RandomF and RandomR (primer sequences are listed in **Supplemental Data S4**) and a guide RNA-encoding template oligonucleotides (guide RNA 20-mer sequences are listed in **Supplemental Data S2**). They were cloned into the BsmBI-linearized blasticidin-resistant lentiCRISPR vector^57^ using Gibson Assembly (New England Biolabs). The resulting lentiCRISPR plasmids were sequenced and named as follows: pAVA2866 (UPF1 KO); pAVA3523 (RPP38 KO); pAVA3545 (RPP29 KO); pAVA3498 (POP5 KO); pAVA3546 (RPP20 KO); pAVA3507 (RPP14 KO); pAVA3573 (RPP30 KO); pAVA3540 (RPP40 KO); pAVA3325 (RPP21 KO); pAVA3547 (RPP24(c18orf21)#1 KO); pAVA3555 (RPP24(c18orf21)#2 KO); pAVA3563 (RPP24(c18orf21)#3 KO); pAVA3500 (RPP64(c3orf17/NEPRO)#1 KO); pAVA3572 (RPP64(c3orf17/NEPRO)#3 KO); pAVA3773 (Control(-)).

#### LentiCRISPR plasmids expressing dual-sgRNAs

DNA fragments containing, in succession, 20-mer sequence of guide RNA#1, Cas9 scaffold, 7SK promoter, and 20-mer sequence of guide RNA#2 were PCR-amplified using primers sgRNA1F and sgRNA2R (**Supplemental Data S4**) that contained sequences of sgRNA#1 and sgRNA#2 (**Supplemental Data S2**), respectively, and plasmid pAVA3129 (**Supplemental Data S3**) as a template. The resulting 332-base-pair DNA fragments were cloned into the BsmBI-linearized blasticidin-resistant lentiCRISPR vector using Gibson Assembly (New England Biolabs) and sequenced. The resulting dual-sgRNA-expressing lentiCRISPR plasmids were sequenced and named as follows: pAVA3584 (RMRP set1 KO); pAVA3583 (RMRP set2 KO); pAVA3586 (RPPH1 set1 KO); pAVA3585 (RPPH1 set2 KO).

#### Plasmids expressing FLAG-tagged proteins

Human RPP64 was PCR-amplified from human cDNA using primers c18_F2 and c18-Flag_R; human RPP24 was PCR-amplified from human cDNA using primers c3_F2 and c3-Flag_R; human RPP21 was PCR-amplified from Addgene plasmid#134542^58^ using primers RPP21_F2 and RPP21-Flag_R; human RPP25 was amplified from Addgene plasmid#134544^58^ using primers RPP25_F2 and RPP25-Flag_R. The resulting PCR products were assembled, together with the C-terminal FLAG-encoding double-stranded DNA fragment formed by annealing oligonucleotides Full-1xFlag-C_F and Full-1xFlag-C_R (**Supplemental Data S4**), into BamHI- and XhoI-digested plasmid pcDNA3 using Gibson Assembly (New England Biolabs). The resulting plasmids were sequenced and named as follows: pAVA3889(CMV-RPP64-FLAG), pAVA3890(CMV-RPP24-FLAG); pAVA3892(CMV-RPP21-FLAG); and pAVA3895(CMV-RPP25-FLAG).

Sequences of the cloning oligonucleotides are listed in **Supplemental Data S4**. Full sequences of the plasmids are shown in **Supplemental Data S3**; they will be deposited to the Addgene repository by the time of publication.

### Construction of the customized lentiviral genome-wide sgRNA omission library

Genome-wide lentiviral sgRNA “omission” library, which deliberately lacked known sgRNAs targeting components and regulators of the nonsense-mediated mRNA degradation (NMD) pathway, contained 90,260 sgRNAs (**Supplemental Data S1**) targeting 20,861 human genes. It was constructed using custom pool of DNA oligonucleotides (**Supplemental Data S1**) ordered from CustomArray, Inc., which were extended using primers RandomF and RandomR (**Supplemental Data S4**) using 17 cycles of PCR (98°C × 3min; 17 cycles of 98°C × 20 sec, 63°C × 30 sec, 72°C × 3 min; 72°C × 10 min; 4°C), which produced a pool of 140-nucleotide DNA products. These DNA products were Gibson-cloned into the BsmBI-linearized blasticidin-resistant lentiCRISPR vector^57^ and transformed into E. cloni 10G electrocompetent cells, producing more than 1000 colonies per each sgRNA in the library. The library of plasmids was isolated directly from plate-grown cells (omitting all steps of growth in liquid culture) using ZymoPure II Plasmid Maxiprep Kit.

### Cell culture and maintenance

The Fireworks reporter cell line^30^ was maintained in DMEM media supplemented with 10% fetal bovine serum (FBS), 100 U/mL penicillin-streptomycin, 150 µg/mL hygromycin, and 0.16 µg/mL puromycin. HEK293T cell line was maintained in DMEM media supplemented with 10% fetal bovine serum (FBS) and 100 U/mL penicillin-streptomycin. All cell lines are authenticated using Short Tandem Repeat (STR) analysis and confirmed to be mycoplasma-free using PCR-based tests.

### Production of lentiviruses and transduction of human cell lines

#### Production of lentiviruses

16 hours prior to the transfection, HEK293T cells were seeded at 60% confluency. They were co-transfected with LentiCRISPR plasmids (or LentiCRISPR omission library) together with the pCMV-dR8.91 packaging and pMD2.G envelope plasmids using the TransIT-293 Transfection Reagent. 24 hours after transfection, the media was changed to DMEM supplemented with 30% fetal bovine serum (FBS) and 100 U/mL penicillin-streptomycin. Lentivirus-containing supernatants were collected at 48 and 72 hours post-transfection and combined. Cellular debris were removed by centrifugation at 200 g for 6 minutes, followed by one passage through a 0.45 µm filter.

#### Transduction of the Fireworks cell line

Fireworks cells were seeded on 15 cm plates at 40% confluency in 20 ml of DMEM supplemented with 10% FBS, 100 U/mL penicillin-streptomycin, and 27 µg/mL polybrene, and transduced using 22 ml of the lentivirus-containing supernatant. 3 days after the infection, cells were selected using 3.0 µg/mL blasticidin for 4 days.

### Genome-wide iterative forward genetic screening

2×10^8^ (8 × 15cm plates) of Fireworks cells were transduced with the Customized Lentiviral Genome-Wide sgRNA Omission Library and propagated in DMEM media supplemented with 10% fetal bovine serum (FBS), 100 U/mL penicillin-streptomycin, 150 µg/mL hygromycin, and 0.16 µg/mL puromycin. 48 hours after infection, blasticidin was added to the media to the final concentration of 3.0 µg/mL for 4 days to select infected cells. Twelve days after infection, cell populations exhibiting right-and-down shift in fluorescence (blue sorting gate in **Fig. 2B**) were FACS-isolated using the Bio-Rad S3e cell sorter. The FACS-isolated cells were pelleted by centrifugation at 1000g for 10 minutes and frozen at -80°C. Genomic DNA was purified from the FACS-isolated cells using phenol extraction. As described earlier^51^, sequences of guide RNAs were amplified from genomic DNA in two steps. First, a linear sgRNA amplification was performed using Herculase II DNA polymerase using 13 thermal cycles with a single sgRNA promoter-specific primer, RandomF (sequences of all primers are listed in **Supplemental Data S4**), and the following cycling parameters: 96°C 20s, 63°C 1min, 72°C 90s. Then, the second PCR primer, RandomR, was added to the reaction and a regular PCR was performed for 35 cycles as follows: 96°C 3min; 35 cycles of 96°C 20s, 63°C 1min, 72°C 45s; 72°C 10min; 4°C. The PCR-amplified pools of sgRNAs were Illumina-sequenced and/or cloned into a BsmBI-linearized, blasticidin-resistant lentiCRISPR vector^57^ to create an enriched lentiCRISPR sgRNA library for subsequent forward genetic screening rounds, as illustrated in **Fig. S2**. After the third enrichment round, the FACS-screening-enriched pool of sgRNAs and the pool of sgRNAs in the original Customized Lentiviral Genome-Wide sgRNA Omission Library used for viral transduction were Illumina-sequenced. For each sgRNA in these pools, the enrichment coefficient was calculated as the ratio of sgRNA abundances after and before the screening. 0.2% of sgRNAs with an extremely low abundance (read count below 10) in the original Customized Lentiviral Genome-Wide sgRNA Omission Library were excluded from ranking (**Supplemental Data S1**). Processing of deep sequencing data was performed as previously described^51^.

### Analysis of CRISPR-generated knockouts

For each knockout, one million cells from the FACS-isolated “Knockout” and “Control(-)” cell populations (**Fig. S3A**) were pelleted by centrifugation and their genomic DNA was phenol-extracted and used as a template to amplify sgRNA-targeted loci using two-step nested PCR (96°C 3 min; 35 cycles of 96°C 30 sec, 63°C 30 sec, 72°C 1 min; 72°C 10 min; 4°C) using the primer pairs listed in **Supplemental Data S4**. The resulting pools of PCR products were subjected to the third round of PCR (98°C 3 min; 5 cycles of 98°C 30 sec, 60°C 30 sec, 72°C 10 min; 72°C 10 min; 4°C) to add Illumina P5 and P7 adaptor sequences using primers P5_sqF and P7_sqR (**Supplemental Data S4**). The pools of adaptor-containing PCR products were sequenced for 110 cycles using NextSeq 500/550 Mid Output Kit according to manufacturer’s instructions with the average depth of 1.4 million reads per sequenced genomic locus. The filtered reads were compared to the wild-type genomic sequences of the respective loci in the GRCh38/hg38 reference human genome. For protein-coding genes, the reads were categorized (**Fig. 3D** and **S3B, C**) as follows: (i) “wild-type” if they either exactly matched the wild-type reference DNA sequence or contained silent mutations resulting in no change in the wild-type protein’s amino acid sequence; (ii) “truncations” if they contained either a stop codon or a frameshift resulting in premature protein termination; (iii) “amino acid mutations” if they caused in-frame amino acid substitutions; and (iv) “uncharacterized splice-site-proximal intronic mutations” if they affected a proximal intronic sequence up to sixteen nucleotides from the splice site. For non-coding genes (RMRP and RPPH1), the reads were categorized (**Fig. 3D** and **S3C**) as follows: “wild-type” if they matched the wild-type reference sequence exactly; (ii) “truncations” if they contained fewer nucleotides than present at the corresponding locus in the reference human genome; (iii) “insertions” if they contained more nucleotides than present at the corresponding locus in the reference human genome; and (iv) “mutations” if they contained nucleotide substitutions not changing the number of nucleotides. The analysis code is deposited at: https://github.com/StoneChen-Clemson/CRISPR-Knockout-analysis.

### Analysis of RNase MRP and RNase P processing

FACS-isolated populations of lentivirus-transduced cells expressing Cas9 and gene-targeting individual sgRNAs: pAVA2866 (UPF1 KO); pAVA3523 (RPP38 KO); pAVA3545 (RPP29 KO); pAVA3498 (POP5 KO); pAVA3546 (RPP20 KO); pAVA3507 (RPP14 KO); pAVA3573 (RPP30 KO); pAVA3540 (RPP40 KO); pAVA3325 (RPP21 KO); pAVA3547 (RPP24(c18orf21)#1 KO); pAVA3555 (RPP24(c18orf21)#2 KO); pAVA3563 (RPP24(c18orf21)#3 KO); pAVA3500 (RPP64(c3orf17/NEPRO)#1 KO); pAVA3572 (RPP64(c3orf17/NEPRO)#3 KO); pAVA3773 (Control(-)), or dual sgRNAs: pAVA3584 (RMRP set1 KO); pAVA3583 (RMRP set2 KO); pAVA3586 (RPPH1 set1 KO); pAVA3585 and (RPPH1 set2 KO) were plated in 6-well plates for 3 hours, washed with phosphate buffered saline, lysed using TRI Reagent, and stored at −80°C. Total RNA was extracted from frozen cells using the TRI Reagent manufacturer’s protocol, treated with DNase I (Promega) for 15 minutes, and purified by phenol extraction and ethanol precipitation. cDNA was synthesized according to the SuperScript IV First-Strand Synthesis System manufacturer’s protocol. The levels of RNase P-unprocessed endogenous lncRNA MALAT1-mascRNA were quantified using nested qPCR across the RNase P cleavage site. Initially, a 20-cycle pre-amplification was performed using Applied Biosystems’ SYBR Green PCR Master Mix with primers JD08-F and JD08-R. The product was then diluted 40-fold and served as the template for qPCR with primers JD07-F and JD07-R. Normalization was performed using 18S rRNA primers H.18SrRNA_F and H.18SrRNA_R. The levels of RNase MRP-unprocessed pre-rRNA at ITS1 site 2 were quantified using RT-qPCR across the site with primers ITS1_2_F1 and ITS_1_2_R2, normalized to 18S rRNA (primers: H.18SrRNA_F and H.18SrRNA_R). Primer sequences are listed in **Supplemental Data S4**.

ΔΔCt was used for quantification of the relative RNA levels. In qPCR bar graphs, the data are presented as means of at least n=3 replicates; error bars represent standard deviation. Statistically significant differences between knockout and control samples were determined by one-way ANOVA. Posthoc comparisons using Tukey’s HSD test were conducted to determine the overall difference between groups, and labeled as “*”, P<0.05; “**”, P<0.01; “***”, P<0.001; “****”, P<0.0001.

### RNA Immunoprecipitation qPCR (RIP-qPCR)

16 hours before transfection, HEK293T cells were seeded on 15cm plates at 40% density in DMEM media supplemented with 10% fetal bovine serum (FBS) and 100 U/mL penicillin-streptomycin. 50 µg of plasmid pAVA3890(CMV-RPP24-FLAG), pAVA3889(CMV-RPP64-FLAG), pAVA3895(CMV-RPP25-FLAG), pAVA3892(CMV-RPP21-FLAG), pAVA1596(CMV-METTL1-FLAG), or pAVA1221(No ORF control) was transfected per plate using the TransIT-293 Transfection Reagent. 24 hours post-transfection, the media was changed to DMEM media supplemented with 10% fetal bovine serum (FBS) and 100 U/mL penicillin-streptomycin. 48 hours post-transfection, cells were washed twice with PBS, pelleted by centrifugation at 200 g for 6 minutes, and placed on ice. Cell pellets were resuspended in 800 µl of pre-chilled buffer A [10 mM Hepes (pH 7.5), 2 mM MgCl_2_, 0.1M KCl, 1 mM DTT] and then passed with eight passages through a 25-gauge needle using a 1ml syringe. Cell nuclei were pelleted by centrifugation at 14,000 g for 30 seconds and then resuspended in 800 µl of buffer B [0.2 M Hepes (pH 7.5), 10% (wt/vol) glycerol, 2 mM MgCl_2_, 0.42 M NaCl, 1 mM DTT, 0.04% NP40, RiboLock RNase Inhibitor, and protease inhibitors cocktail]. The lysates were rotated at 4°C for 1 hour and then centrifuged at 14,000 g for 5 minutes. 650 µl of the supernatant representing clarified nuclear extract was transferred to a new tube. For each sample, 50 µl of the clarified nuclear extract were combined with 1 ml of TRI Reagent and stored at −80°C for subsequent RNA purification as input controls. For immunoprecipitations, 30 µl of anti-Flag M2 affinity gel beads was resuspended in 200 µl of buffer B, combined with 500 µl of the clarified nuclear extract, rotated overnight at 4°C, and washed ten times with 800 µl of buffer B, each time pelleting the beads at 5000 g for 1 minute and transferring the beads to a new tube. FLAG-tagged proteins were eluted by addition of 100 µl of 1 µg/ul 3xFLAG peptide resuspended in Buffer B to 30 µl of washed beads for one hour. The mixture was briefly centrifuged at 5000 g for one minute and then 80 µl of the FLAG-tagged protein-containing supernatant were combined with 1 ml of TRI Reagent and stored at −80°C for subsequent RNA isolation. RNA was isolated according to the TRI Reagent manufacturer’s protocol, treated with DNase I (Promega) for 15 minutes, and purified by phenol extraction and ethanol precipitation. cDNA was synthesized according to the SuperScript IV First-Strand Synthesis System manufacturer’s protocol. qPCR quantification of RPPH1 RNA was performed using primers H1_F1 and H1_R1, and RMRP RNA using primers RMRP_qF1 and RMRP_qR1. ΔΔCt was used for quantification of the relative RNA levels. The relative enrichment of RPPH1 RNA and RMRP RNA in the FLAG-purified fractions was quantified by normalizing their RNA levels to those in the inputs, and then referencing them to the enrichment of RPPH1 RNA and RMRP RNA in the FLAG-purified fraction of the “No ORF” negative control. The sequences of primers used in this study are listed in **Supplemental Data S4**.

