## Supplemental Figures and Legends for "Identification of Two Elusive Human Ribonuclease MRP-Specific Protein Components"

**Figure S1. Application of the Fireworks reporter system to detect RNase MRP/P deficiency via inhibition of the nonsense mediated decay caused by a translation defect, related to Figure 1. A.** Schematic of the previously described Fireworks reporter system[30]. Each Fireworks reporter expresses a single polyprotein consisting of (i) fluorescent proteins (GFP or RFP), (ii) TEV protease, and (iii) PEST-β-globin. The β-globin in the RFP (but not in GFP) reporter carries a premature termination codon (PTC39) upstream of its last (second) intron. The resulting polyprotein contains TEV protease, which separates translated fluorescent proteins, protease, and the PEST-β-globin via in vivo proteolytic cleavage. The PEST degradation sequence was added to shorten β-globin half-life. **B.** Schematics of fluorescence response of the Fireworks reporters[30] shown in panel A to inhibition of nonsense-mediated mRNA decay (NMD), translation, and transcription. **C.** CRISPR knockouts of the human RNase MRP- and RNase P-specific RNA genes, RMRP and RPPH1, using additional guide RNAs reproduce right-and-down fluorescence shift in the Fireworks cell line.

**Figure S2. Schematic of the iterative rounds of guide RNA library-based forward genetic screening, related to Figure 2.** Details of each step are described in Methods Details.

**Figure S3. Generation and analysis of individual knockouts of screening-identified genes, related to Figure 3. A.** Schematic of generating Knockout and Control(-) cell populations for downstream analysis. Populations of right-and-down shifted cells in the blue FACS gate were collected and lysed with Trizol right after their attachment to plates, ensuring efficient collection of cells with gene knockout. **B.** Individual FACS-based validation of the screening-identified RPP24 and RPP64 using additional sgRNAs and deep sequencing analysis of their knockouts. **C.** Deep-sequencing analysis of genomic sites analyzed in **Fig. 3D** in cells transduced with a non-targeting negative control sgRNA.

**Figure S4. Sequence alignment of human c3orf17 (RPP64) and c18orf21 (RPP24) with proteins in other organisms, related to Figure 4. A.** Alignment of *H. sapiens* ENSG00000163608 (Ensembl) with *X. laevis* XB-GENE-5928625 (Xenbase), *D. melanogaster* FBgn0036036 (Flybase), *S. japonicum* SJAG_03952 (JaponicusDB), *S. pombe* SPAC323.08

(PomBase), and *S. cerevisiae* YLR145W (Saccharomyces Genome Database). Amino acid sequences of cryo-EM-determined (PDB: 7C7A[21]) alpha-helices αI-αIV in yeast *S. cerevisiae* Rmp1 (YLR145W) are underlined. **B.** Alignment of *H. sapiens* ENSG00000141428 (Ensembl) with *X. laevis* XB-GENE-995475 (Xenbase), *D. melanogaster* FBgn0053082 (Flybase), *S. japonicum* SJAG_02547 (JaponicusDB), *S. pombe* SPCC16C4.19 (PomBase), and *S. cerevisiae* YDR478W (Saccharomyces Genome Database). Amino acid sequences of two cryo-EM-determined (PDB: 7C7A[21]) zinc-coordinating CXXC motifs in yeast *S. cerevisiae* Snm1 (YDR478W) are indicated with red boxes. Multiple sequence alignments were performed using Clustal Omega[55] and formatted using MView[56].

### Figure S1

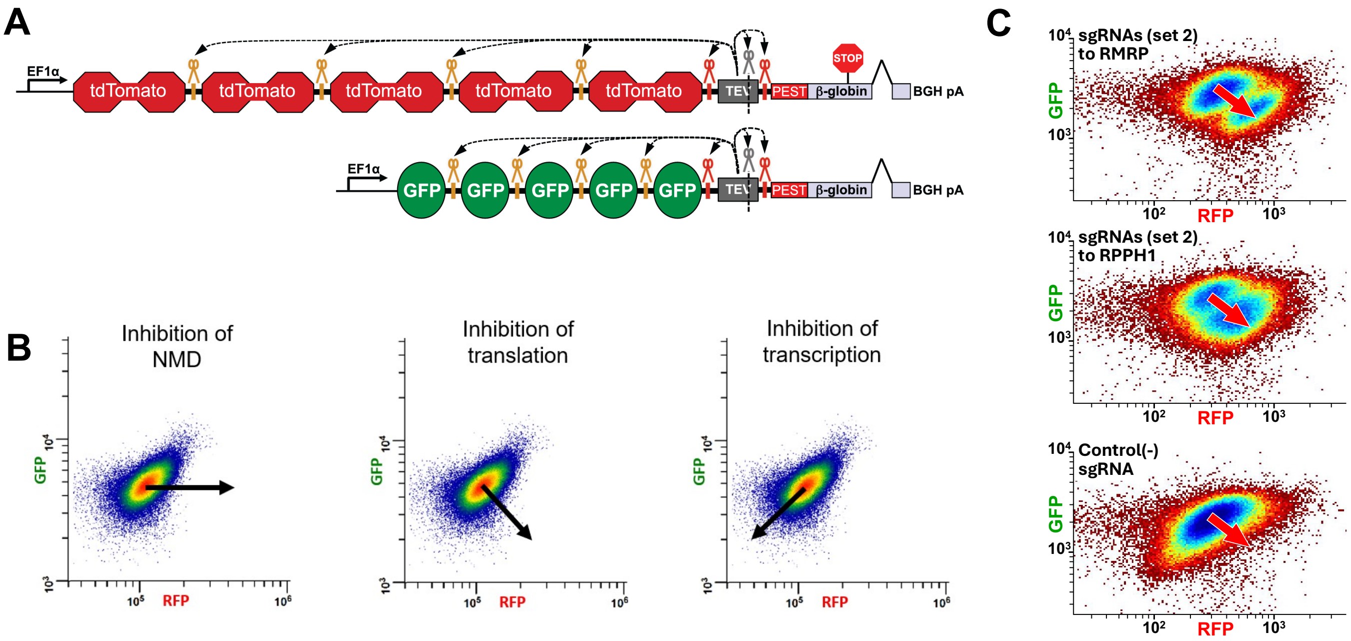
A
EF1α
tdTomato
tdTomato
tdTomato
tdTomato
tdTomato
TEV
PEST
β-globin
STOP
BGH pA
EF1α
GFP
GFP
GFP
GFP
GFP
TEV
PEST
β-globin
BGH pA
B
Inhibition of NMD
Inhibition of translation
Inhibition of transcription
GFP
RFP
C
sgRNAs (set 2) to RMRP
sgRNAs (set 2) to RPPH1
Control(-) sgRNA
GFP
RFP

### Figure S2

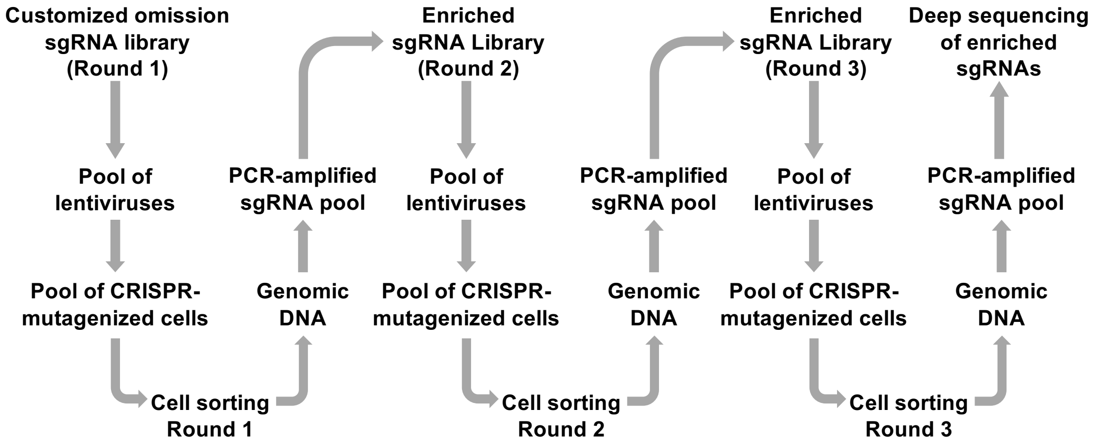
Customized omission sgRNA library (Round 1)
Pool of lentiviruses
Pool of CRISPR-mutagenized cells
Cell sorting Round 1
Genomic DNA
PCR-amplified sgRNA pool
Enriched sgRNA Library (Round 2)
Pool of lentiviruses
Pool of CRISPR-mutagenized cells
Cell sorting Round 2
Genomic DNA
PCR-amplified sgRNA pool
Enriched sgRNA Library (Round 3)
Pool of lentiviruses
Pool of CRISPR-mutagenized cells
Cell sorting Round 3
Genomic DNA
PCR-amplified sgRNA pool
Deep sequencing of enriched sgRNAs

### Figure S3

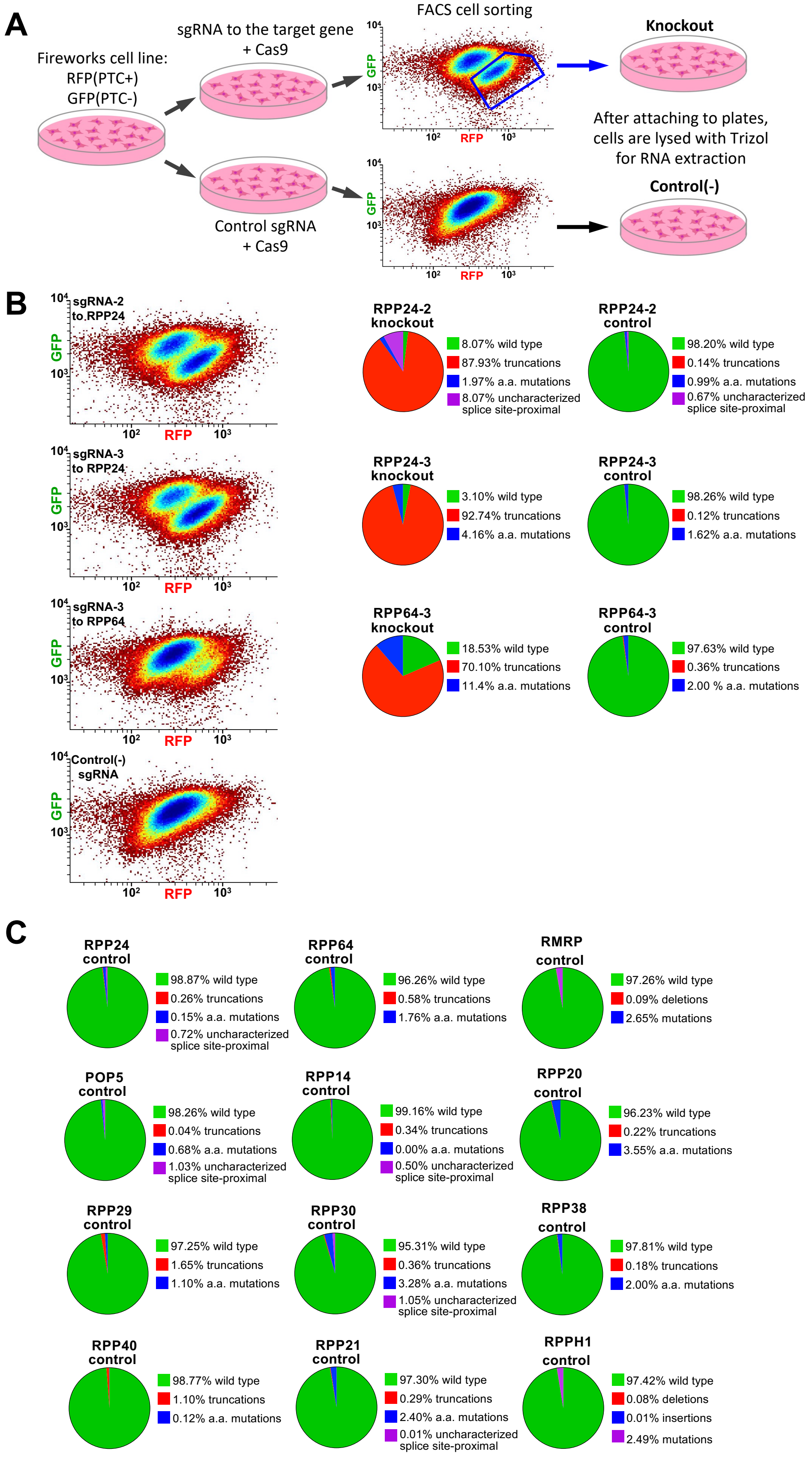
A
Fireworks cell line:
RFP(PTC+)
GFP(PTC-)
sgRNA to the target gene
+ Cas9
Control sgRNA
+ Cas9
FACS cell sorting
Knockout
After attaching to plates, cells are lysed with Trizol for RNA extraction
Control(-)
B
sgRNA-2 to RPP24
sgRNA-3 to RPP24
sgRNA-3 to RPP64
Control(-) sgRNA
GFP
RFP
RPP24-2 knockout
8.07% wild type
87.93% truncations
1.97% a.a. mutations
8.07% uncharacterized splice site-proximal
RPP24-2 control
98.20% wild type
0.14% truncations
0.99% a.a. mutations
0.67% uncharacterized splice site-proximal
RPP24-3 knockout
3.10% wild type
92.74% truncations
4.16% a.a. mutations
RPP24-3 control
98.26% wild type
0.12% truncations
1.62% a.a. mutations
RPP64-3 knockout
18.53% wild type
70.10% truncations
11.4% a.a. mutations
RPP64-3 control
97.63% wild type
0.36% truncations
2.00 % a.a. mutations
C
RPP24 control
98.87% wild type
0.26% truncations
0.15% a.a. mutations
0.72% uncharacterized splice site-proximal
RPP64 control
96.26% wild type
0.58% truncations
1.76% a.a. mutations
RMRP control
97.26% wild type
0.09% deletions
2.65% mutations
POP5 control
98.26% wild type
0.04% truncations
0.68% a.a. mutations
1.03% uncharacterized splice site-proximal
RPP14 control
99.16% wild type
0.34% truncations
0.00% a.a. mutations
0.50% uncharacterized splice site-proximal
RPP20 control
96.23% wild type
0.22% truncations
3.55% a.a. mutations
RPP29 control
97.25% wild type
1.65% truncations
1.10% a.a. mutations
RPP30 control
95.31% wild type
0.36% truncations
3.28% a.a. mutations
1.05% uncharacterized splice site-proximal
RPP38 control
97.81% wild type
0.18% truncations
2.00% a.a. mutations
RPP40 control
98.77% wild type
1.10% truncations
0.12% a.a. mutations
RPP21 control
97.30% wild type
0.29% truncations
2.40% a.a. mutations
0.01% uncharacterized splice site-proximal
RPPH1 control
97.42% wild type
0.08% deletions
0.01% insertions
2.49% mutations

### Figure S4

**A**

#### c3orf17 (RPP64)

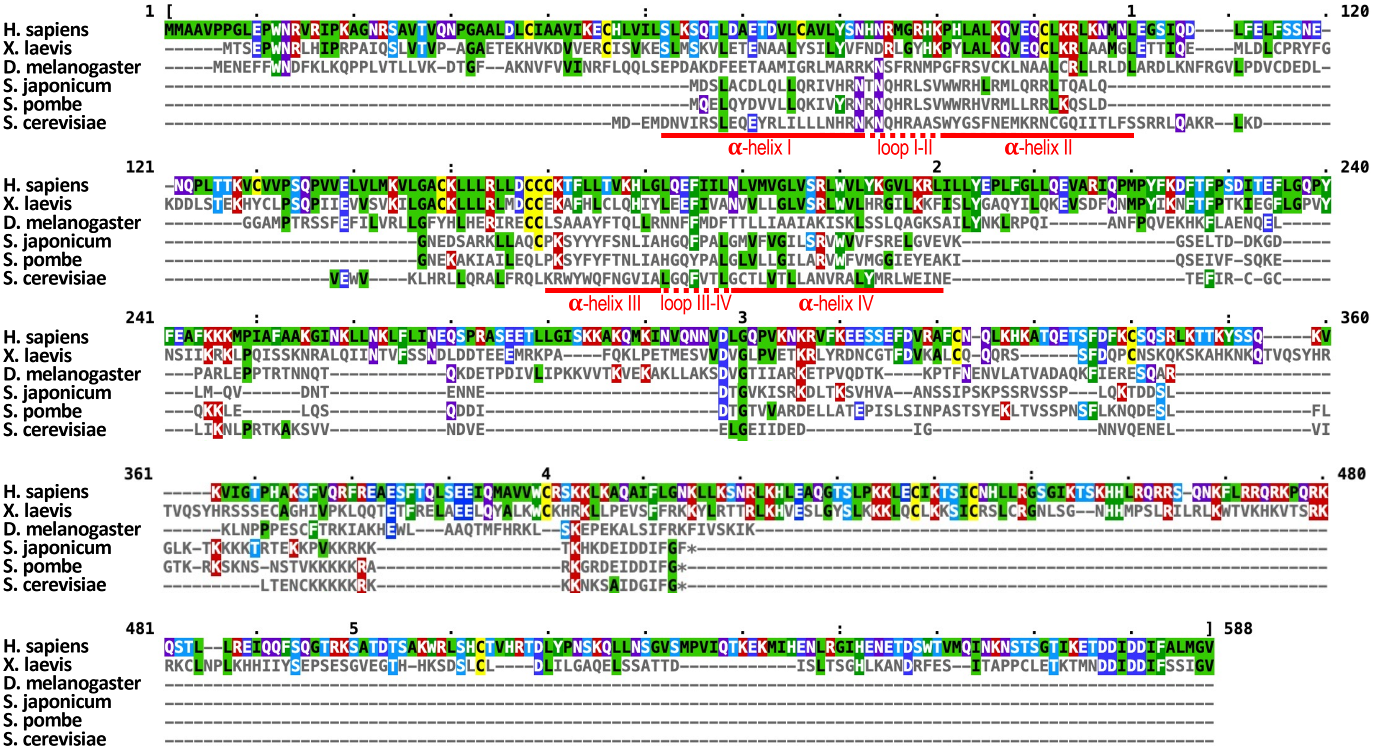


**B**

#### c18orf21 (RPP24)

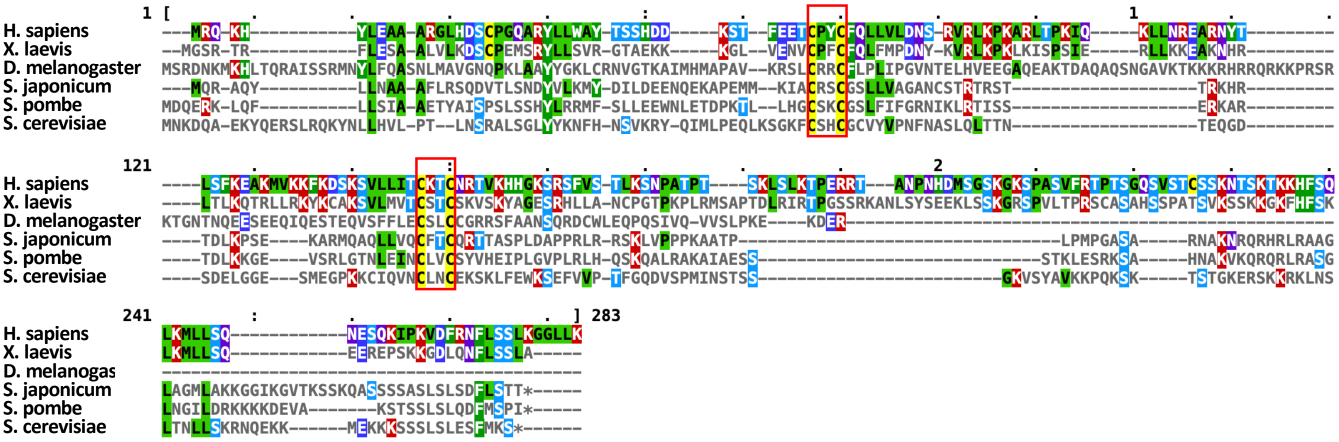
