## Supplemental Data Descriptions for "Identification of Two Elusive Human Ribonuclease MRP-Specific Protein Components"

### DESCRIPTION OF SUPPLEMENTAL DATA

**Supplemental Data S1.** Guide RNA omission library and guide RNA enrichment by iterative rounds of forward genetic screening.

**Supplemental Data S2.** Sequences of guide RNAs for CRISPR knockouts.

**Supplemental Data S3.** Plasmid sequences.

**Supplemental Data S4.** Sequences of oligonucleotides.
