## Supplemental Data S3 for "Identification of Two Elusive Human Ribonuclease MRP-Specific Protein Components"

### Supplemental Data File S1: Supplemental vector sequences, related to Experimental Procedures

#### A) pAVA3889 [CMV-RPP64-FLAG]:

**CMV promotor: 235-818**
**RPP64: 916-2616**
**FLAG: 2629-2652**
**BGH ployA: 2738-2962**

gacggatcgggagatctcccgatcccctatggtcgactctcagtacaatctgctctgatgccgc
atagttaagccagtatctgctccctgcttgtgtgttggaggtcgctgagtagtgcgcgagcaaa
atttaagctacaacaaggcaaggcttgaccgacaattgcatgaagaatctgcttagggttaggc
gttttgcgctgcttcgcgatgtacgggccagatatacgcgttgacattgattattgactagtta
ttaatagtaatcaattacggggtcattagttcatagcccatatatggagttccgcgttacataa
cttacggtaaatggcccgcctggctgaccgcccaacgacccccgcccattgacgtcaataatga
cgtatgttcccatagtaacgccaatagggactttccattgacgtcaatgggtggactatttacg
gtaaactgcccacttggcagtacatcaagtgtatcatatgccaagtacgccccctattgacgtc
aatgacggtaaatggcccgcctggcattatgcccagtacatgaccttatgggactttcctactt
ggcagtacatctacgtattagtcatcgctattaccatggtgatgcggttttggcagtacatcaa
tgggcgtggatagcggtttgactcacggggatttccaagtctccaccccattgacgtcaatggg
agtttgttttggcaccaaaatcaacgggactttccaaaatgtcgtaacaactccgccccattga
cgcaaatgggcggtaggcgtgtacggtgggaggtctatataagcagagctctctggctaactag
agaacccactgcttactggcttatcgaaattaatacgactcactatagggagacccaagcttgg
taccgagctcGGATCCaccATGATGGCTGCGGTGCCGCCGGGCCTGGAGCCGTGGAACCGTGTG
AGAATCCCTAAGGCGGGGAACCGCAGCGCAGTGACAGTGCAGAACCCCGGCGCGGCCCTTGACC
TTTGCATTGCAGCTGTAATTAAAGAATGCCATCTCGTCATACTGTCGCTGAAGAGCCAAACCTT
AGATGCAGAAACAGATGTGTTATGTGCAGTCCTTTACAGCAATCACAACAGAATGGGCCGCCAC
AAACCCCATTTGGCCCTCAAACAGGTTGAGCAATGTTTAAAGCGTTTGAAAAACATGAATTTGG
AGGGCTCAATTCAAGACCTGTTTGAGTTGTTTTCTTCCAATGAAAATCAGCCCTTAACTACCAA
AGTATGTGTTGTCCCCAGTCAGCCAGTGGTGGAGTTGGTGTTGATGAAGGTTTTGGGAGCCTGC
AAGTTGTTGCTCCGCTTGTTGGACTGCTGCTGCAAAACTTTTCTTTTGACTGTGAAACATCTAG
GTTTGCAAGAGTTCATTATTTTAAACCTTGTGATGGTTGGGCTGGTGAGCAGGTTATGGGTTCT
CTATAAAGGTGTCTTAAAAAGGTTGATTTTGTTATATGAGCCTTTGTTTGGATTGCTTCAAGAG
GTCGCTAGGATTCAACCAATGCCTTACTTCAAAGATTTTACCTTTCCTTCTGATATCACTGAAT
TTTTAGGACAGCCATATTTTGAAGCCTTTAAGAAAAAAATGCCTATAGCTTTTGCAGCTAAAGG
AATAAATAAATTGCTAAATAAACTGTTTTTAATAAATGAGCAGTCACCAAGAGCCAGTGAAGAA
ACCTTGCTTGGAATTTCAAAAAAAGCTAAACAAATGAAGATCAATGTACAGAATAATGTGGATC
TTGGACAGCCAGTAAAGAATAAGAGAGTCTTCAAAGAAGAGTCATCAGAATTTGATGTGAGGGC
TTTCTGCAACCAGCTGAAACACAAAGCTACTCAGGAGACCAGTTTTGATTTTAAATGTTCTCAA

TCCAGACTAAAGACAACCAAGTATTCTTCTCAGAAAGTGATAGGAACTCCTCATGCCAAAAGTT
TTGTGCAAAGATTCCGAGAGGCTGAGTCCTTCACACAACTTTCTGAAGAAATCCAGATGGCAGT
TGTATGGTGCAGGAGCAAAAAACTCAAGGCTCAGGCCATTTTTCTGGGTAACAAACTTCTTAAA
AGCAACCGGCTTAAACATCTGGAAGCTCAAGGTACTAGTTTGCCAAAGAAACTAGAGTGCATAA
AAACGTCTATTTGCAACCACCTTCTTCGTGGCTCAGGTATCAAAACTTCAAAGCATCATCTGAG
ACAGAGAAGATCACAGAATAAATTTTTACGGAGACAAAGGAAACCACAGAGAAAGTTGCAGTCG
ACTCTTTTAAGGGAAATTCAGCAGTTCTCTCAAGGGACTCGGAAGAGTGCTACAGATACCAGTG
CTAAGTGGAGACTCTCACACTGTACTGTGCATAGAACTGATCTCTACCCTAACAGTAAGCAGCT
CTTGAATAGTGGAGTTTCAATGCCTGTCATACAAACTAAGGAGAAAATGATTCATGAAAATCTT
AGAGGCATCCATGAAAATGAAACTGATTCGTGGACGGTGATGCAAATAAATAAAAACAGTACAT
CAGGAACCATTAAGGAGACAGATGACATTGATGATATTTTTGCTTTAATGGGAGTTatcgatgt
cgaggattacaaggatgacgacgataagtagggcccggtaccCTCGAGcatgcatctagagggc
cctattctatagtgtcacctaaatgctagagctcgctgatcagcctcgactgtgccttctagtt
gccagccatctgttgtttgcccctcccccgtgccttccttgaccctggaaggtgccactcccac
tgtcctttcctaataaaatgaggaaattgcatcgcattgtctgagtaggtgtcattctattctg
gggggtggggtggggcaggacagcaagggggaggattgggaagacaatagcaggcatgctgggg
atgcggtgggctctatggcttctgaggcggaaagaaccagctggggctctagggggtatcccca
cgcgccctgtagcggcgcattaagcgcggcgggtgtggtggttacgcgcagcgtgaccgctaca
cttgccagcgccctagcgcccgctcctttcgctttcttcccttcctttctcgccacgttcgccg
gctttccccgtcaagctctaaatcgggggcatccctttagggttccgatttagtgctttacggca
cctcgaccccaaaaaacttgattagggtgatggttcacgtagtgggccatcgccctgatagacg
gtttttcgccctttgacgttggagtccacgttctttaatagtggactcttgttccaaactggaa
caacactcaaccctatctcggtctattcttttgatttataagggattttggggatttcggccta
ttggttaaaaaatgagctgatttaacaaaaatttaacgcgaattaattctgtggaatgtgtgtc
agttagggtgtggaaagtccccaggctccccaggcaggcagaagtatgcaaagcatgcatctca
attagtcagcaaccaggtgtggaaagtccccaggctccccagcaggcagaagtatgcaaagcat
gcatctcaattagtcagcaaccatagtcccgcccctaactccgcccatcccgcccctaactccg
cccagttccgcccattctccgccccatggctgactaattttttttatttatgcagaggccgagg
ccgcctctgcctctgagctattccagaagtagtgaggaggcttttttggaggcctaggcttttg
caaaaagctcccgggagcttgtatatccattttcggatctgatcaagagacaggatgaggatcg
tttcgcatgattgaacaagatggattgcacgcaggttctccggccgcttgggtggagaggctat
tcggctatgactgggcacaacagacaatcggctgctctgatgccgccgtgttccggctgtcagc
gcaggggcgcccggttctttttgtcaagaccgacctgtccggtgccctgaatgaactgcaggac
gaggcagcgcggctatcgtggctggccacgacgggcgttccttgcgcagctgtgctcgacgttg
tcactgaagcgggaagggactggctgctattgggcgaagtgccggggcaggatctcctgtcatc
tcaccttgctcctgccgagaaagtatccatcatggctgatgcaatgcggcggctgcatacgctt
gatccggctacctgcccattcgaccaccaagcgaaacatcgcatcgagcgagcacgtactcgga
tggaagccggtcttgtcgatcaggatgatctggacgaagagcatcaggggctcgcgccagccga
actgttcgccaggctcaaggcgcgcatgcccgacggcgaggatctcgtcgtgacccatggcgat
gcctgcttgccgaatatcatggtggaaaatggccgcttttctggattcatcgactgtggccggc
tgggtgtggcggaccgctatcaggacatagcgttggctacccgtgatattgctgaagagcttgg

cggcgaatgggctgaccgcttcctcgtgctttacggtatcgccgctcccgattcgcagcgcatc
gccttctatcgccttcttgacgagttcttctgagcgggactctggggttcgaaatgaccgacca
agcgacgcccaacctgccatcacgagatttcgattccaccgccgccttctatgaaaggttgggc
ttcggaatcgttttccgggacgccggctggatgatcctccagcgcggggatctcatgctggagt
tcttcgcccaccccaacttgtttattgcagcttataatggttacaaataaagcaatagcatcac
aaatttcacaaataaagcatttttttcactgcattctagttgtggtttgtccaaactcatcaat
gtatcttatcatgtctgtataccgtcgacctctagctagagcttggcgtaatcatggtcatagc
tgtttcctgtgtgaaattgttatccgctcacaattccacacaacatacgagccggaagcataaa
gtgtaaagcctggggtgcctaatgagtgagctaactcacattaattgcgttgcgctcactgccc
gctttccagtcgggaaacctgtcgtgccagctgcattaatgaatcggccaacgcgcggggagag
gcggtttgcgtattgggcgctcttccgcttcctcgctcactgactcgctgcgctcggtcgttcg
gctgcggcgagcggtatcagctcactcaaaggcggtaatacggttatccacagaatcaggggat
aacgcaggaaagaacatgtgagcaaaaggccagcaaaaggccaggaaccgtaaaaaggccgcgt
tgctggcgtttttccataggctccgcccccctgacgagcatcacaaaaatcgacgctcaagtca
gaggtggcgaaacccgacaggactataaagataccaggcgtttccccctggaagctccctcgtg
cgctctcctgttccgaccctgccgcttaccggatacctgtccgcctttctcccttcgggaagcg
tggcgctttctcaatgctcacgctgtaggtatctcagttcggtgtaggtcgttcgctccaagct
gggctgtgtgcacgaaccccccgttcagcccgaccgctgcgccttatccggtaactatcgtctt
gagtccaacccggtaagacacgacttatcgccactggcagcagccactggtaacaggattagca
gagcgaggtatgtaggcggtgctacagagttcttgaagtggtggcctaactacggctacactag
aaggacagtatttggtatctgcgctctgctgaagccagttaccttcggaaaaagagttggtagc
tcttgatccggcaaacaaaccaccgctggtagcggtggtttttttgtttgcaagcagcagatta
cgcgcagaaaaaaaggatctcaagaagatcctttgatcttttctacggggtctgacgctcagtg
gaacgaaaactcacgttaagggattttggtcatgagattatcaaaaaggatcttcacctagatc
cttttaaattaaaaatgaagttttaaatcaatctaaagtatatatgagtaaacttggtctgaca
gttaccaatgcttaatcagtgaggcacctatctcagcgatctgtctatttcgttcatccatagt
tgcctgactccccgtcgtgtagataactacgatacgggagggcttaccatctggccccagtgct
gcaatgataccgcgagacccacgctcaccggctccagatttatcagcaataaaccagccagccg
gaagggccgagcgcagaagtggtcctgcaactttatccgcctccatccagtctattaattgttg
ccgggaagctagagtaagtagttcgccagttaatagtttgcgcaacgttgttgccattgctaca
ggcatcgtggtgtcacgctcgtcgtttggtatggcttcattcagctccggttcccaacgatcaa
ggcgagttacatgatcccccatgttgtgcaaaaaagcggttagctccttcggtcctccgatcgt
tgtcagaagtaagttggccgcagtgttatcactcatggttatggcagcactgcataattctctt
actgtcatgccatccgtaagatgcttttctgtgactggtgagtactcaaccaagtcattctgag
aatagtgtatgcggcgaccgagttgctcttgcccggcgtcaatacgggataataccgcgccaca
tagcagaactttaaaagtgctcatcattggaaaacgttcttcggggcgaaaactctcaaggatc
ttaccgctgttgagatccagttcgatgtaacccactcgtgcacccaactgatcttcagcatctt
ttactttcaccagcgtttctgggtgagcaaaaacaggaaggcaaaatgccgcaaaaaagggaat
aagggcgacacggaaatgttgaatactcatactcttcctttttcaatattattgaagcatttat
cagggttattgtctcatgagcggatacatatttgaatgtatttagaaaaataaacaaataggggg
ttccgcgcacatttccccgaaaagtgccacctgacgtc

**B) pAVA3890 [CMV-RPP24-FLAG]:**

**CMV promotor: 235-818**
**RPP64: 961-1575**
**FLAG: 1588-1611**
**BGH ployA: 1697-1921**

```
gacggatcgggagatctcccgatcccctatggtcgactctcagtacaatctgctctgatgccgc
atagttaagccagtatctgctccctgcttgtgtgttggaggtcgctgagtagtgcgcgagcaaa
atttaagctacaacaaggcaaggcttgaccgacaattgcatgaagaatctgcttagggttaggc
gttttgcgctgcttcgcgatgtacgggccagatatacgcgttgacattgattattgactagtta
ttaatagtaatcaattacggggtcattagttcatagcccatatatggagttccgcgttacataa
cttacggtaaatggcccgcctggctgaccgcccaacgacccccgcccattgacgtcaataatga
cgtatgttcccatagtaacgccaatagggactttccattgacgtcaatgggtggactatttacg
gtaaactgcccacttggcagtacatcaagtgtatcatatgccaagtacgccccctattgacgtc
aatgacggtaaatggcccgcctggcattatgcccagtacatgaccttatgggactttcctactt
ggcagtacatctacgtattagtcatcgctattaccatggtgatgcggttttggcagtacatcaa
tgggcgtggatagcggtttgactcacggggatttccaagtctccaccccattgacgtcaatggg
agtttgttttggcaccaaaatcaacgggactttccaaaatgtcgtaacaactccgccccattga
cgcaaatgggcggtaggcgtgtacggtgggaggtctatataagcagagctctctggctaactag
agaacccactgcttactggcttatcgaaattaatacgactcactatagggagacccaagcttgg
taccgagctcGGATCCaccATGAGACAGAAGCACTACCTTGAGGCTGCAGCGCGGGACTGCAC
GACAGCTGCCCGGGCCAAGCCCGCTACCTCCTCTGGGCCTACACTTCGTCGCACGATGATAAGA
GCACTTTTGAAGAAACGTGTCCATACTGTTTCCAGCTGTTGGTTCTGGATAACTCTCGAGTGCG
TCTCAAACCCAAAGCCAGGTTGACACCCAAAATACAGAAACTTCTTAATCGAGAAGCGAGAAAC
TATACACTCAGTTTTAAAGAAGCAAAAATGGTGAAAAAGTTCAAAGACTCCAAAAGTGTATTGT
TGATCACTTGTAAAACATGCAACAGAACAGTGAAACATCATGGTAAAAGTAGAAGCTTTGTGTC
AACATTGAAGAGCAATCCTGCCACTCCTACAAGTAAACTCAGCCTGAAGACACCAGAGAGAAGG
ACTGCAAACCCAAATCATGACATGTCTGGCTCGAAAGGCAAGAGCCCAGCATCGGTTTTCAGAA
CACCTACATCTGGACAGTCAGTATCTACTTGCTCCTCAAAGAACACCAGCAAAACAAAGAAACA
CTTCTCTCAACTAAAAATGTTACTTAGTCAGAATGAATCCCAAAAGATTCCAAAGGTGGACTTC
AGAAATTTCTTATCTTCTCTGAAGGGTGGACTTTTAAAAatcgatgtcgaggattacaaggatg
acgacgataagtagggcccggtaccCTCGAGcatgcatctagagggccctattctatagtgtca
cctaaatgctagagctcgctgatcagcctcgactgtgccttctagttgccagccatctgttgtt
tgcccctcccccgtgccttccttgaccctggaaggtgccactcccactgtcctttcctaataaa
atgaggaaattgcatcgcattgtctgagtaggtgtcattctattctggggggtggggtggggca
ggacagcaagggggaggattgggaagacaatagcaggcatgctggggatgcggtgggctctatg
gcttctgaggcggaaagaaccagctggggctctagggggtatccccacgcgccctgtagcggcg
cattaagcgcggcgggtgtggtggttacgcgcagcgtgaccgctacacttgccagcgccctagc
gcccgctcctttcgctttcttcccttcctttctcgccacgttcgccggctttccccgtcaagct
ctaaatcggggcatccctttagggttccgatttagtgctttacggcacctcgaccccaaaaaac
```

ttgattagggtgatggttcacgtagtgggccatcgccctgatagacggtttttcgccctttgac
gttggagtccacgttctttaatagtggactcttgttccaaactggaacaacactcaaccctatc
tcggtctattcttttgatttataagggattttggggatttcggcctattggttaaaaaatgagc
tgatttaacaaaaatttaacgcgaattaattctgtggaatgtgtgtcagttagggtgtggaaag
tccccaggctccccaggcaggcagaagtatgcaaagcatgcatctcaattagtcagcaaccagg
tgtggaaagtccccaggctccccagcaggcagaagtatgcaaagcatgcatctcaattagtcag
caaccatagtcccgcccctaactccgcccatcccgcccctaactccgcccagttccgcccattc
tccgccccatggctgactaattttttttatttatgcagaggccgaggccgcctctgcctctgag
ctattccagaagtagtgaggaggcttttttggaggcctaggcttttgcaaaaagctcccgggag
cttgtatatccattttcggatctgatcaagagacaggatgaggatcgtttcgcatgattgaaca
agatggattgcacgcaggttctccggccgcttgggtggagaggctattcggctatgactgggca
caacagacaatcggctgctctgatgccgccgtgttccggctgtcagcgcaggggcgcccggttc
tttttgtcaagaccgacctgtccggtgccctgaatgaactgcaggacgaggcagcgcggctatc
gtggctggccacgacgggcgttccttgcgcagctgtgctcgacgttgtcactgaagcgggaagg
gactggctgctattgggcgaagtgccggggcaggatctcctgtcatctcaccttgctcctgccg
agaaagtatccatcatggctgatgcaatgcggcggctgcatacgcttgatccggctacctgccc
attcgaccaccaagcgaaacatcgcatcgagcgagcacgtactcggatggaagccggtcttgtc
gatcaggatgatctggacgaagagcatcaggggctcgcgccagccgaactgttcgccaggctca
aggcgcgcatgcccgacggcgaggatctcgtcgtgacccatggcgatgcctgcttgccgaatat
catggtggaaaatggccgcttttctggattcatcgactgtggccggctgggtgtggcggaccgc
tatcaggacatagcgttggctacccgtgatattgctgaagagcttggcggcgaatgggctgacc
gcttcctcgtgctttacggtatcgccgctcccgattcgcagcgcatcgccttctatcgccttct
tgacgagttcttctgagcgggactctggggttcgaaatgaccgaccaagcgacgcccaacctgc
catcacgagatttcgattccaccgccgccttctatgaaaggttgggcttcggaatcgttttccg
ggacgccggctggatgatcctccagcgcggggatctcatgctggagttcttcgcccaccccaac
ttgtttattgcagcttataatggttacaaataaagcaatagcatcacaaatttcacaaataaag
catttttttcactgcattctagttgtggtttgtccaaactcatcaatgtatcttatcatgtctg
tataccgtcgacctctagctagagcttggcgtaatcatggtcatagctgtttcctgtgtgaaat
tgttatccgctcacaattccacacaacatacgagccggaagcataaagtgtaaagcctggggtg
cctaatgagtgagctaactcacattaattgcgttgcgctcactgcccgctttccagtcgggaaa
cctgtcgtgccagctgcattaatgaatcggccaacgcgcggggagaggcggtttgcgtattggg
cgctcttccgcttcctcgctcactgactcgctgcgctcggtcgttcggctgcggcgagcggtat
cagctcactcaaaggcggtaatacggttatccacagaatcaggggataacgcaggaaagaacat
gtgagcaaaaggccagcaaaaggccaggaaccgtaaaaaggccgcgttgctggcgtttttccat
aggctccgcccccctgacgagcatcacaaaaatcgacgctcaagtcagaggtggcgaaacccga
caggactataaagataccaggcgtttccccctggaagctccctcgtgcgctctcctgttccgac
cctgccgcttaccggatacctgtccgcctttctcccttcgggaagcgtggcgctttctcaatgc
tcacgctgtaggtatctcagttcggtgtaggtcgttcgctccaagctgggctgtgtgcacgaac
cccccgttcagcccgaccgctgcgccttatccggtaactatcgtcttgagtccaacccggtaag
acacgacttatcgccactggcagcagccactggtaacaggattagcagagcgaggtatgtaggc
ggtgctacagagttcttgaagtggtggcctaactacggctacactagaaggacagtatttggta

tctgcgctctgctgaagccagttaccttcggaaaaagagttggtagctcttgatccggcaaaca
aaccaccgctggtagcggtggtttttttgtttgcaagcagcagattacgcgcagaaaaaaagga
tctcaagaagatcctttgatcttttctacggggtctgacgctcagtggaacgaaaactcacgtt
aagggattttggtcatgagattatcaaaaaggatcttcacctagatccttttaaattaaaaatg
aagttttaaatcaatctaaagtatatatgagtaaacttggtctgacagttaccaatgcttaatc
agtgaggcacctatctcagcgatctgtctatttcgttcatccatagttgcctgactccccgtcg
tgtagataactacgatacgggagggcttaccatctggccccagtgctgcaatgataccgcgaga
cccacgctcaccggctccagatttatcagcaataaaccagccagccggaagggccgagcgcaga
agtggtcctgcaactttatccgcctccatccagtctattaattgttgccgggaagctagagtaa
gtagttcgccagttaatagtttgcgcaacgttgttgccattgctacaggcatcgtggtgtcacg
ctcgtcgtttggtatggcttcattcagctccggttcccaacgatcaaggcgagttacatgatcc
cccatgttgtgcaaaaaagcggttagctccttcggtcctccgatcgttgtcagaagtaagttgg
ccgcagtgttatcactcatggttatggcagcactgcataattctcttactgtcatgccatccgt
aagatgcttttctgtgactggtgagtactcaaccaagtcattctgagaatagtgtatgcggcga
ccgagttgctcttgcccggcgtcaatacgggataataccgcgccacatagcagaactttaaaag
tgctcatcattggaaaacgttcttcggggcgaaaactctcaaggatcttaccgctgttgagatc
cagttcgatgtaacccactcgtgcacccaactgatcttcagcatcttttactttcaccagcgtt
tctgggtgagcaaaaacaggaaggcaaaatgccgcaaaaaagggaataagggcgacacggaaat
gttgaatactcatactcttcctttttcaatattattgaagcatttatcagggttattgtctcat
gagcggatacatatttgaatgtatttagaaaaataaacaaataggggttccgcgcacatttccc
cgaaaagtgccacctgacgtc

**C) pAVA3892 [CMV-RPP21-FLAG]**

**CMV promotor: 235-818**
**RPP21: 961-1377**
**FLAG: 1390-1413**
**BGH polyA: 1499-1723**

gacggatcgggagatctcccgatcccctatggtcgactctcagtacaatctgctctgatgccgc
atagttaagccagtatctgctccctgcttgtgtgttggaggtcgctgagtagtgcgcgagcaaa
atttaagctacaacaaggcaaggcttgaccgacaattgcatgaagaatctgcttagggttaggc
gttttgcgctgcttcgcgatgtacgggccagatatacgcgttgacattgattattgactagtta
ttaatagtaatcaattacggggtcattagttcatagcccatatatggagttccgcgttacataa
cttacggtaaatggcccgcctggctgaccgcccaacgacccccgcccattgacgtcaataatga
cgtatgttcccatagtaacgccaatagggactttccattgacgtcaatgggtggactatttacg
gtaaactgcccacttggcagtacatcaagtgtatcatatgccaagtacgccccctattgacgtc
aatgacggtaaatggcccgcctggcattatgcccagtacatgaccttatgggactttcctactt
ggcagtacatctacgtattagtcatcgctattaccatggtgatgcggttttggcagtacatcaa
tgggcgtggatagcggtttgactcacggggatttccaagtctccaccccattgacgtcaatggg

agtttgttttggcaccaaaatcaacgggactttccaaaatgtcgtaacaactccgccccattga
cgcaaatgggcggtaggcgtgtacggtgggaggtctatataagcagagctctctggctaactag
agaacccactgcttactggcttatcgaaattaatacgactcactatagggagacccaagcttgg
taccgagctcGGATCCaccATGGCGGGGCCGGTGAAGGACCGCGAGGCCTTCCAGAGGCTCAAC
TTCCTGTACCAGGCCGCCCATTGTGTCCTTGCCCAGGACCCCGAGAACCAGGCGCTGGCGAGGT
TTTACTGCTACACTGAGAGGACCATTGCGAAGCGGCTCGTCTTGCGGCGAGATCCCTCGGTGAA
GAGGACTCTCTGTCGAGGCTGCTCTTCCCTCCTCGTCCCGGGCCTCACCTGCACCCAGCGCCAG
AGACGCTGCAGGGGACAGCGCTGGACCGTACAGACCTGCCTAACATGCCAGCGCAGCCAACGCT
TCCTCAATGATCCCGGGCATTTACTCTGGGGAGACAGGCCTGAGGCCCAGCTCGGGAGCCAAGC
AGATTCCAAACCACTACAACCCTTGCCAAACACAGCCCACTCCATTTCAGACCGCCTTCCTGAG
GAGAAAATGCAGACTCAGGGTTCCAGTAACCAGatcgatgtcgaggattacaaggatgacgacg
ataagtagggcccggtaccCTCGAGcatgcatctagagggccctattctatagtgtcacctaaa
tgctagagctcgctgatcagcctcgactgtgccttctagttgccagccatctgttgtttgcccc
tcccccgtgccttccttgaccctggaaggtgccactcccactgtcctttcctaataaaatgagg
aaattgcatcgcattgtctgagtaggtgtcattctattctggggggtggggtggggcaggacag
caagggggaggattgggaagacaatagcaggcatgctggggatgcggtgggctctatggcttct
gaggcggaaagaaccagctggggctctagggggtatccccacgcgccctgtagcggcgcattaa
gcgcggcgggtgtggtggttacgcgcagcgtgaccgctacacttgccagcgccctagcgcccgc
tcctttcgctttcttcccttcctttctcgccacgttcgccggctttccccgtcaagctctaaat
cgggggcatccctttagggttccgatttagtgctttacggcacctcgaccccaaaaaacttgatt
agggtgatggttcacgtagtgggccatcgccctgatagacggtttttcgccctttgacgttgga
gtccacgttctttaatagtggactcttgttccaaactggaacaacactcaaccctatctcggtc
tattcttttgatttataagggattttggggatttcggcctattggttaaaaaatgagctgattt
aacaaaaatttaacgcgaattaattctgtggaatgtgtgtcagttagggtgtggaaagtcccca
ggctccccaggcaggcagaagtatgcaaagcatgcatctcaattagtcagcaaccaggtgtgga
aagtccccaggctccccagcaggcagaagtatgcaaagcatgcatctcaattagtcagcaacca
tagtcccgcccctaactccgcccatcccgcccctaactccgcccagttccgcccattctccgcc
ccatggctgactaattttttttatttatgcagaggccgaggccgcctctgcctctgagctattc
cagaagtagtgaggaggcttttttggaggcctaggcttttgcaaaaagctcccgggagcttgta
tatccattttcggatctgatcaagagacaggatgaggatcgtttcgcatgattgaacaagatgg
attgcacgcaggttctccggccgcttgggtggagaggctattcggctatgactgggcacaacag
acaatcggctgctctgatgccgccgtgttccggctgtcagcgcaggggcgcccggttctttttg
tcaagaccgacctgtccggtgccctgaatgaactgcaggacgaggcagcgcggctatcgtggct
ggccacgacgggcgttccttgcgcagctgtgctcgacgttgtcactgaagcgggaagggactgg
ctgctattgggcgaagtgccggggcaggatctcctgtcatctcaccttgctcctgccgagaaag
tatccatcatggctgatgcaatgcggcggctgcatacgcttgatccggctacctgcccattcga
ccaccaagcgaaacatcgcatcgagcgagcacgtactcggatggaagccggtcttgtcgatcag
gatgatctggacgaagagcatcaggggctcgcgccagccgaactgttcgccaggctcaaggcgc
gcatgcccgacggcgaggatctcgtcgtgacccatggcgatgcctgcttgccgaatatcatggt
ggaaaatggccgcttttctggattcatcgactgtggccggctgggtgtggcggaccgctatcag
gacatagcgttggctacccgtgatattgctgaagagcttggcggcgaatggctgaccgcttcc

tcgtgctttacggtatcgccgctcccgattcgcagcgcatcgccttctatcgccttcttgacga
gttcttctgagcgggactctggggttcgaaatgaccgaccaagcgacgcccaacctgccatcac
gagatttcgattccaccgccgccttctatgaaaggttgggcttcggaatcgttttccgggacgc
cggctggatgatcctccagcgcggggatctcatgctggagttcttcgcccaccccaacttgttt
attgcagcttataatggttacaaataaagcaatagcatcacaaatttcacaaataaagcatttt
tttcactgcattctagttgtggtttgtccaaactcatcaatgtatcttatcatgtctgtatacc
gtcgacctctagctagagcttggcgtaatcatggtcatagctgtttcctgtgtgaaattgttat
ccgctcacaattccacacaacatacgagccggaagcataaagtgtaaagcctggggtgcctaat
gagtgagctaactcacattaattgcgttgcgctcactgcccgctttccagtcgggaaacctgtc
gtgccagctgcattaatgaatcggccaacgcgcggggagaggcggtttgcgtattgggcgctct
tccgcttcctcgctcactgactcgctgcgctcggtcgttcggctgcggcgagcggtatcagctc
actcaaaggcggtaatacggttatccacagaatcaggggataacgcaggaaagaacatgtgagc
aaaaggccagcaaaaggccaggaaccgtaaaaaggccgcgttgctggcgtttttccataggctc
cgcccccctgacgagcatcacaaaaatcgacgctcaagtcagaggtggcgaaacccgacaggac
tataaagataccaggcgtttccccctggaagctccctcgtgcgctctcctgttccgaccctgcc
gcttaccggatacctgtccgcctttctcccttcgggaagcgtggcgctttctcaatgctcacgc
tgtaggtatctcagttcggtgtaggtcgttcgctccaagctgggctgtgtgcacgaaccccccg
ttcagcccgaccgctgcgccttatccggtaactatcgtcttgagtccaacccggtaagacacga
cttatcgccactggcagcagccactggtaacaggattagcagagcgaggtatgtaggcggtgct
acagagttcttgaagtggtggcctaactacggctacactagaaggacagtatttggtatctgcg
ctctgctgaagccagttaccttcggaaaaagagttggtagctcttgatccggcaaacaaaccac
cgctggtagcggtggtttttttgtttgcaagcagcagattacgcgcagaaaaaaaggatctcaa
gaagatcctttgatcttttctacggggtctgacgctcagtggaacgaaaactcacgttaaggga
ttttggtcatgagattatcaaaaaggatcttcacctagatccttttaaattaaaaatgaagttt
taaatcaatctaaagtatatatgagtaaacttggtctgacagttaccaatgcttaatcagtgag
gcacctatctcagcgatctgtctatttcgttcatccatagttgcctgactccccgtcgtgtaga
taactacgatacgggagggcttaccatctggccccagtgctgcaatgataccgcgagacccacg
ctcaccggctccagatttatcagcaataaaccagccagccggaagggccgagcgcagaagtggt
cctgcaactttatccgcctccatccagtctattaattgttgccgggaagctagagtaagtagtt
cgccagttaatagtttgcgcaacgttgttgccattgctacaggcatcgtggtgtcacgctcgtc
gtttggtatggcttcattcagctccggttcccaacgatcaaggcgagttacatgatcccccatg
ttgtgcaaaaaagcggttagctccttcggtcctccgatcgttgtcagaagtaagttggccgcag
tgttatcactcatggttatggcagcactgcataattctcttactgtcatgccatccgtaagatg
cttttctgtgactggtgagtactcaaccaagtcattctgagaatagtgtatgcggcgaccgagt
tgctcttgcccggcgtcaatacgggataataccgcgccacatagcagaactttaaaagtgctca
tcattggaaaacgttcttcggggcgaaaactctcaaggatcttaccgctgttgagatccagttc
gatgtaacccactcgtgcacccaactgatcttcagcatcttttactttcaccagcgtttctggg
tgagcaaaaacaggaaggcaaaatgccgcaaaaaagggaataagggcgacacggaaatgttgaa
tactcatactcttcctttttcaatattattgaagcatttatcagggttattgtctcatgagcgg
atacatatttgaatgtatttagaaaaataaacaaataggggttccgcgcacatttccccgaaaa
gtgccacctgacgtc

**D) pAVA3895 [CMV-RPP25-FLAG]**

**CMV promotor: 235-818**
**RPP25: 961-1512**
**FLAG: 1525-1548**
**BGH polyA: 1634-1858**

gacggatcgggagatctcccgatcccctatggtcgactctcagtacaatctgctctgatgccgc
atagttaagccagtatctgctccctgcttgtgtgttggaggtcgctgagtagtgcgcgagcaaa
atttaagctacaacaaggcaaggcttgaccgacaattgcatgaagaatctgcttagggttaggc
gttttgcgctgcttcgcgatgtacgggccagatatacgcgttgacattgattattgactagtta
ttaatagtaatcaattacggggtcattagttcatagcccatatatggagttccgcgttacataa
cttacggtaaatggcccgcctggctgaccgcccaacgacccccgcccattgacgtcaataatga
cgtatgttcccatagtaacgccaatagggactttccattgacgtcaatgggtggactatttacg
gtaaactgcccacttggcagtacatcaagtgtatcatatgccaagtacgccccctattgacgtc
aatgacggtaaatggcccgcctggcattatgcccagtacatgaccttatgggactttcctactt
ggcagtacatctacgtattagtcatcgctattaccatggtgatgcggttttggcagtacatcaa
tgggcgtggatagcggtttgactcacggggatttccaagtctccaccccattgacgtcaatggg
agtttgttttggcaccaaaatcaacgggactttccaaaatgtcgtaacaactccgccccattga
cgcaaatgggcggtaggcgtgtacggtgggaggtctatataagcagagctctctggctaactag
agaacccactgcttactggcttatcgaaattaatacgactcactatagggagacccaagcttgg
taccgagctcGGATCCaccATGGAGAACTTCCGTAAGGTGCGCTCCGAAGAGGCGCCAGCGGGG
TGCGGGGCCGAGGGAGGCGGCCCGGGCTCCGGCCCCTTCGCAGACCTGGCGCCGGGCGCGGTGC
ACATGCGGGTCAAGGAAGGCAGCAAGATCCGGAACCTGATGGCCTTCGCCACCGCCAGCATGGC
GCAGCCAGCCACGCGCGCCATCGTCTTCAGCGGCTGCGGCCGGGCCACCACCAAAACCGTCACG
TGCGCCGAGATCCTCAAGCGCCGCCTGGCGGGCCTGCACCAGGTCACGCGGCTGCGCTACCGGA
GCGTACGCGAGGTGTGGCAGAGCCTCCCGCCTGGGCCCACGCAGGGTCAGACGCCTGGCGAGCC
GGCCGCTAGTCTCAGCGTACTTAAGAACGTGCCCGGCCTCGCCATCCTACTTTCCAAGGACGCG
CTGGATCCGCGACAGCCCGGCTACCAGCCCCCGAATCCCCATCCTGGTCCCTCGTCCCCGCCAG
CCGCGCCAGCGTCCAAGAGGAGCCTAGGGGAACCCGCAGCTGGAGAAGGCTCCGCGAAGCGATC
GCAACCCGAGCCAGGGGTTGCGGACGAGGATCAGACGGCCatcgatgtcgaggattacaaggat
gacgacgataagtagggcccggtaccCTCGAGcatgcatctagagggccctattctatagtgtc
acctaaatgctagagctcgctgatcagcctcgactgtgccttctagttgccagccatctgttgt
ttgcccctcccccgtgccttccttgaccctggaaggtgccactcccactgtcctttcctaataa
aatgaggaaattgcatcgcattgtctgagtaggtgtcattctattctggggggtggggtggggc
aggacagcaagggggaggattgggaagacaatagcaggcatgctggggatgcggtgggctctat
ggcttctgaggcggaaagaaccagctggggctctagggggtatccccacgcgccctgtagcggc
gcattaagcgcggcgggtgtggtggttacgcgcagcgtgaccgctacacttgccagcgccctag

cgcccgctcctttcgctttcttcccttcctttctcgccacgttcgccggctttccccgtcaagc
tctaaatcggggcatccctttagggttccgatttagtgctttacggcacctcgaccccaaaaaa
cttgattagggtgatggttcacgtagtgggccatcgccctgatagacggtttttcgccctttga
cgttggagtccacgttctttaatagtggactcttgttccaaactggaacaacactcaaccctat
ctcggtctattcttttgatttataagggattttggggatttcggcctattggttaaaaaatgag
ctgatttaacaaaaatttaacgcgaattaattctgtggaatgtgtgtcagttagggtgtggaaa
gtccccaggctccccaggcaggcagaagtatgcaaagcatgcatctcaattagtcagcaaccag
gtgtggaaagtccccaggctccccagcaggcagaagtatgcaaagcatgcatctcaattagtca
gcaaccatagtcccgcccctaactccgcccatcccgcccctaactccgcccagttccgcccatt
ctccgccccatggctgactaattttttttatttatgcagaggccgaggccgcctctgcctctga
gctattccagaagtagtgaggaggcttttttggaggcctaggcttttgcaaaaagctcccggga
gcttgtatatccattttcggatctgatcaagagacaggatgaggatcgtttcgcatgattgaac
aagatggattgcacgcaggttctccggccgcttgggtggagaggctattcggctatgactgggc
acaacagacaatcggctgctctgatgccgccgtgttccggctgtcagcgcaggggcgcccggtt
ctttttgtcaagaccgacctgtccggtgccctgaatgaactgcaggacgaggcagcgcggctat
cgtggctggccacgacgggcgttccttgcgcagctgtgctcgacgttgtcactgaagcgggaag
ggactggctgctattgggcgaagtgccggggcaggatctcctgtcatctcaccttgctcctgcc
gagaaagtatccatcatggctgatgcaatgcggcggctgcatacgcttgatccggctacctgcc
cattcgaccaccaagcgaaacatcgcatcgagcgagcacgtactcggatggaagccggtcttgt
cgatcaggatgatctggacgaagagcatcaggggctcgcgccagccgaactgttcgccaggctc
aaggcgcgcatgcccgacggcgaggatctcgtcgtgacccatggcgatgcctgcttgccgaata
tcatggtggaaaatggccgcttttctggattcatcgactgtggccggctgggtgtggcggaccg
ctatcaggacatagcgttggctacccgtgatattgctgaagagcttggcggcgaatgggctgac
cgcttcctcgtgctttacggtatcgccgctcccgattcgcagcgcatcgccttctatcgccttc
ttgacgagttcttctgagcgggactctggggttcgaaatgaccgaccaagcgacgcccaacctg
ccatcacgagatttcgattccaccgccgccttctatgaaaggttgggcttcggaatcgttttcc
gggacgccggctggatgatcctccagcgcggggatctcatgctggagttcttcgcccaccccaa
cttgtttattgcagcttataatggttacaaataaagcaatagcatcacaaatttcacaaataaa
gcatttttttcactgcattctagttgtggtttgtccaaactcatcaatgtatcttatcatgtct
gtataccgtcgacctctagctagagcttggcgtaatcatggtcatagctgtttcctgtgtgaaa
ttgttatccgctcacaattccacacaacatacgagccggaagcataaagtgtaaagcctggggt
gcctaatgagtgagctaactcacattaattgcgttgcgctcactgcccgctttccagtcgggaa
acctgtcgtgccagctgcattaatgaatcggccaacgcgcggggagaggcggtttgcgtattgg
gcgctcttccgcttcctcgctcactgactcgctgcgctcggtcgttcggctgcggcgagcggta
tcagctcactcaaaggcggtaatacggttatccacagaatcaggggataacgcaggaaagaaca
tgtgagcaaaaggccagcaaaaggccaggaaccgtaaaaaggccgcgttgctggcgtttttcca
taggctccgcccccctgacgagcatcacaaaaatcgacgctcaagtcagaggtggcgaaacccg
acaggactataaagataccaggcgtttccccctggaagctccctcgtgcgctctcctgttccga
ccctgccgcttaccggatacctgtccgcctttctcccttcgggaagcgtggcgctttctcaatg
ctcacgctgtaggtatctcagttcggtgtaggtcgttcgctccaagctgggctgtgtgcacgaa
ccccccgttcagcccgaccgctgcgccttatccggtaactatcgtcttgagtccaacccggtaa

gacacgacttatcgccactggcagcagccactggtaacaggattagcagagcgaggtatgtagg
cggtgctacagagttcttgaagtggtggcctaactacggctacactagaaggacagtatttggt
atctgcgctctgctgaagccagttaccttcggaaaaagagttggtagctcttgatccggcaaac
aaaccaccgctggtagcggtggtttttttgtttgcaagcagcagattacgcgcagaaaaaaagg
atctcaagaagatcctttgatcttttctacggggtctgacgctcagtggaacgaaaactcacgt
taagggattttggtcatgagattatcaaaaaggatcttcacctagatccttttaaattaaaaat
gaagttttaaatcaatctaaagtatatatgagtaaacttggtctgacagttaccaatgcttaat
cagtgaggcacctatctcagcgatctgtctatttcgttcatccatagttgcctgactccccgtc
gtgtagataactacgatacgggagggcttaccatctggccccagtgctgcaatgataccgcgag
acccacgctcaccggctccagatttatcagcaataaaccagccagccggaagggccgagcgcag
aagtggtcctgcaactttatccgcctccatccagtctattaattgttgccgggaagctagagta
agtagttcgccagttaatagtttgcgcaacgttgttgccattgctacaggcatcgtggtgtcac
gctcgtcgtttggtatggcttcattcagctccggttcccaacgatcaaggcgagttacatgatc
ccccatgttgtgcaaaaaagcggttagctccttcggtcctccgatcgttgtcagaagtaagttg
gccgcagtgttatcactcatggttatggcagcactgcataattctcttactgtcatgccatccg
taagatgcttttctgtgactggtgagtactcaaccaagtcattctgagaatagtgtatgcggcg
accgagttgctcttgcccggcgtcaatacgggataataccgcgccacatagcagaactttaaaa
gtgctcatcattggaaaacgttcttcggggcgaaaactctcaaggatcttaccgctgttgagat
ccagttcgatgtaacccactcgtgcacccaactgatcttcagcatcttttactttcaccagcgt
ttctgggtgagcaaaaacaggaaggcaaaatgccgcaaaaaagggaataagggcgacacggaaa
tgttgaatactcatactcttcctttttcaatattattgaagcatttatcagggttattgtctca
tgagcggatacatatttgaatgtatttagaaaaataaacaaataggggttccgcgcacatttcc
ccgaaaagtgccacctgacgtc

**E) pAVA3129 [lentiviral expression of Cas9 and two sgRNAs: U6-sgRNA1, 7SK-sgRNA2]**

**U6 promoter: 1716-1956**
**sgRNA1: 1967-1986**
**sgRNA Scaffold: 1987-2062**
**7SK promoter: 2075- 2316**
**sgRNA2: 2319-2338**
**sgRNA Scaffold: 2339-2414**
**Filler Sequence (HIV-1 isolate): 2462-2579**
**EF1alfa promoter: 2623-2873**
**Cas9: 2912-7111**
**P2A peptide: 7118-7183**
**Blasticidin$^{R}$: 7184-7582**

TTAATGTAGTCTTATGCAATACTCTTGTAGTCTTGCAACATGGTAACGATGAGTTAGCAACATGCCTTACAAGGAGA
GAAAAAGCACCGTGCATGCCGATTGGTGGAAGTAAGGTGGTACGATCGTGCCTTATTAGGAAGGCAACAGACGGGTC
TGACATGGATTGGACGAACCACTGAATTGCCGCATTGCAGAGATATTGTATTTAAGTGCCTAGCTCGATACATAAAC

GGGTCTCTCTGGTTAGACCAGATCTGAGCCTGGGAGCTCTCTGGCTAACTAGGGAACCCACTGCTTAAGCCTCAATA
AAGCTTGCCTTGAGTGCTTCAAGTAGTGTGTGCCCGTCTGTTGTGTGACTCTGGTAACTAGAGATCCCTCAGACCCT
TTTAGTCAGTGTGGAAAATCTCTAGCAGTGGCGCCCGAACAGGGACTTGAAAGCGAAAGGGAAACCAGAGGAGCTCT
CTCGACGCAGGACTCGGCTTGCTGAAGCGCGCACGGCAAGAGGCGAGGGGCGGCGACTGGTGAGTACGCCAAAAATT
TTGACTAGCGGAGGCTAGAAGGAGAGAGATGGGTGCGAGAGCGTCAGTATTAAGCGGGGGAGAATTAGATCGCGATG
GGAAAAAATTCGGTTAAGGCCAGGGGGAAAGAAAAAATATAAATTAAAACATATAGTATGGGCAAGCAGGGAGCTAG
AACGATTCGCAGTTAATCCTGGCCTGTTAGAAACATCAGAAGGCTGTAGACAAATACTGGGACAGCTACAACCATCC
CTTCAGACAGGATCAGAAGAACTTAGATCATTATATAATACAGTAGCAACCCTCTATTGTGTGCATCAAAGGATAGA
GATAAAAGACACCAAGGAAGCTTTAGACAAGATAGAGGAAGAGCAAAACAAAAGTAAGACCACCGCACAGCAAGCGG
CCGCTGATCTTCAGACCTGGAGGAGGAGATATGAGGGACAATTGGAGAAGTGAATTATATAAATATAAAGTAGTAAA
AATTGAACCATTAGGAGTAGCACCCACCAAGGCAAAGAGAAGAGTGGTGCAGAGAGAAAAAAGAGCAGTGGGAATAG
GAGCTTTGTTCCTTGGGTTCTTGGGAGCAGCAGGAAGCACTATGGGCGCAGCGTCAATGACGCTGACGGTACAGGCC
AGACAATTATTGTCTGGTATAGTGCAGCAGCAGAACAATTTGCTGAGGGCTATTGAGGCGCAACAGCATCTGTTGCA
ACTCACAGTCTGGGGCATCAAGCAGCTCCAGGCAAGAATCCTGGCTGTGGAAAGATACCTAAAGGATCAACAGCTCC
TGGGGATTTGGGGTTGCTCTGGAAAACTCATTTGCACCACTGCTGTGCCTTGGAATGCTAGTTGGAGTAATAAATCT
CTGGAACAGATTTGGAATCACACGACCTGGATGGAGTGGGACAGAGAAATTAACAATTACACAAGCTTAATACACTC
CTTAATTGAAGAATCGCAAAACCAGCAAGAAAAGAATGAACAAGAATTATTGGAATTAGATAAATGGGCAAGTTTGT
GGAATTGGTTTAACATAACAAATTGGCTGTGGTATATAAAATTATTCATAATGATAGTAGGAGGCTTGGTAGGTTTA
AGAATAGTTTTTGCTGTACTTTCTATAGTGAATAGAGTTAGGCAGGGATATTCACCATTATCGTTTCAGACCCACCT
CCCAACCCCGAGGGGACCCAG**AGAGGGCCTATTTCCCATGATTCCTTCATATTTGCATATACGATACAAGGCTGTTA**
**GAGAGATAATTAGAATTAATTTGACTGTAAACACAAAGATATTAGTACAAAATACGTGACGTAGAAAGTAATAATTT**
**CTTGGGTAGTTTGCAGTTTTAAAATTATGTTTTAAAATGGACTATCATATGCTTACCGTAACTTGAAAGTATTTCGA**
**TTTCTTGGCTTTATATATCTTGTGGAAAGGA**CGAAACACC**GNNNNNNNNNNNNNNNNNNNN**GTTTTAGAGCTAGAAA
TAGCAAGTTAAAATAAGGCTAGTCCGTTATCAACTTGAAAAAGTGGCACCGAGTCGGTGCTTTTTTGAATTCctgca
gtatttagcatgccccacccatctgcaaggcattctggatagtgtcaaaacagccggaaatcaagtccgtttatctc
aaactttagcattttgggaataaatgatatttgctatgctggttaaattagattttagttaaatttcctgctgaagc
tctagtacgataagcaacttgacctaagtgtaaagttgagacttccttcaggtt**tatata**gcttgtgcgccgcttgg
gtacct**CGNNNNNNNNNNNNNNNNNNNN**GTTTTAGAGCTAGAAATAGCAAGTTAAAATAAGGCTAGTCCGTTATCAA
CTTGAAAAAGTGGCACCGAGTCGGTGCTTTTTTGAATTCTAGATCTTGAGACAAATGGCAGTATTCATCCACAA**TTT**
**TAAAAGAAAAGGGGGGATTGGGGGGTACAGTGCAGGGGAAAGAATAGTAGACATAATAGCAACAGACATACAAACTA**
**AAGAATTACAAAAACAAATTACAAAAATTCAAAATTTT**CGGGTTTATTACAGGGACAGCAGAGATCCACTTTGGCGC
CGGCTCGAGTGGCTCCGGTGCCCGTCAGTGGGCAGAGCGCACATCGCCCACAGTCCCCGAGAAGTTGGGGGGAGGGG
TCGGCAATTGAACCGGTGCCTAGAGAAGGTGGCGCGGGGTAAACTGGGAAAGTGATGTCGTGTACTGGCTCCGCCTT
TTTCCCGAGGGTGGGGGAGAACCGTATATAAGTGCAGTAGTCGCCGTGAACGTTCTTTTTCGCAACGGGTTTGCCGC
CAGAACACAGGTGTCGTGACGCGGGATCCGCCACCATGGATTACAAAGACGATGACGATAAGATGGCCCCAAAGAAG
AAGCGGAAGGTCGGTATCCACGGAGTCCCAGCAGCCGACAAGAAGTACAGCATCGGCCTGGACATCGGCACCAACTC
TGTGGGCTGGGCCGTGATCACCGACGAGTACAAGGTGCCCAGCAAGAAATTCAAGGTGCTGGGCAACACCGACCGGC
ACAGCATCAAGAAGAACCTGATCGGAGCCCTGCTGTTCGACAGCGGCGAAACAGCCGAGGCCACCCGGCTGAAGAGA
ACCGCCAGAAGAAGATACACCAGACGGAAGAACCGGATCTGCTATCTGCAAGAGATCTTCAGCAACGAGATGGCCAA
GGTGGACGACAGCTTCTTCCACAGACTGGAAGAGTCCTTCCTGGTGGAAGAGGATAAGAAGCACGAGCGGCACCCCA
TCTTCGGCAACATCGTGGACGAGGTGGCCTACCACGAGAAGTACCCCACCATCTACCACCTGAGAAAGAAACTGGTG
GACAGCACCGACAAGGCCGACCTGCGGCTGATCTATCTGGCCCTGGCCCACATGATCAAGTTCCGGGGCCACTTCCT
GATCGAGGGCGACCTGAACCCCGACAACAGCGACGTGGACAAGCTGTTCATCCAGCTGGTGCAGACCTACAACCAGC
TGTTCGAGGAAAACCCCATCAACGCCAGCGGCGTGGACGCCAAGGCCATCCTGTCTGCCAGACTGAGCAAGAGCAGA
CGGCTGGAAAATCTGATCGCCCAGCTGCCCGGCGAGAAGAAGAATGGCCTGTTCGGCAACCTGATTGCCCTGAGCCT
GGGCCTGACCCCCAACTTCAAGAGCAACTTCGACCTGGCCGAGGATGCCAAACTGCAGCTGAGCAAGGACACCTACG
ACGACGACCTGGACAACCTGCTGGCCCAGATCGGCGACCAGTACGCCGACCTGTTTCTGGCCGCCAAGAACCTGTCC
GACGCCATCCTGCTGAGCGACATCCTGAGAGTGAACACCGAGATCACCAAGGCCCCCCTGAGCGCCTCTATGATCAA
GAGATACGACGAGCACCACCAGGACCTGACCCTGCTGAAAGCTCTCGTGCGGCAGCAGCTGCCTGAGAAGTACAAAG

AGATTTTCTTCGACCAGAGCAAGAACGGCTACGCCGGCTACATTGACGGCGGAGCCAGCCAGGAAGAGTTCTACAAG
TTCATCAAGCCCATCCTGGAAAAGATGGACGGCACCGAGGAACTGCTCGTGAAGCTGAACAGAGAGGACCTGCTGCG
GAAGCAGCGGACCTTCGACAACGGCAGCATCCCCCACCAGATCCACCTGGGAGAGCTGCACGCCATTCTGCGGCGGC
AGGAAGATTTTTACCCATTCCTGAAGGACAACCGGGAAAAGATCGAGAAGATCCTGACCTTCCGCATCCCCTACTAC
GTGGGCCCTCTGGCCAGGGGAAACAGCAGATTCGCCTGGATGACCAGAAAGAGCGAGGAAACCATCACCCCCTGGAA
CTTCGAGGAAGTGGTGGACAAGGGCGCTTCCGCCCAGAGCTTCATCGAGCGGATGACCAACTTCGATAAGAACCTGC
CCAACGAGAAGGTGCTGCCCAAGCACAGCCTGCTGTACGAGTACTTCACCGTGTATAACGAGCTGACCAAAGTGAAA
TACGTGACCGAGGGAATGAGAAAGCCCGCCTTCCTGAGCGGCGAGCAGAAAAAGGCCATCGTGGACCTGCTGTTCAA
GACCAACCGGAAAGTGACCGTGAAGCAGCTGAAAGAGGACTACTTCAAGAAAATCGAGTGCTTCGACTCCGTGGAAA
TCTCCGGCGTGGAAGATCGGTTCAACGCCTCCCTGGGCACATACCACGATCTGCTGAAAATTATCAAGGACAAGGAC
TTCCTGGACAATGAGGAAAACGAGGACATTCTGGAAGATATCGTGCTGACCCTGACACTGTTTGAGGACAGAGAGAT
GATCGAGGAACGGCTGAAAACCTATGCCCACCTGTTCGACGACAAAGTGATGAAGCAGCTGAAGCGGCGGAGATACA
CCGGCTGGGGCAGGCTGAGCCGGAAGCTGATCAACGGCATCCGGGACAAGCAGTCCGGCAAGACAATCCTGGATTTC
CTGAAGTCCGACGGCTTCGCCAACAGAAACTTCATGCAGCTGATCCACGACGACAGCCTGACCTTTAAAGAGGACAT
CCAGAAAGCCCAGGTGTCCGGCCAGGGCGATAGCCTGCACGAGCACATTGCCAATCTGGCCGGCAGCCCCGCCATTA
AGAAGGGCATCCTGCAGACAGTGAAGGTGGTGGACGAGCTCGTGAAAGTGATGGGCCGGCACAAGCCCGAGAACATC
GTGATCGAAATGGCCAGAGAGAACCAGACCACCCAGAAGGGACAGAAGAACAGCCGCGAGAGAATGAAGCGGATCGA
AGAGGGCATCAAAGAGCTGGGCAGCCAGATCCTGAAAGAACACCCCGTGGAAAACACCCAGCTGCAGAACGAGAAGC
TGTACCTGTACTACCTGCAGAATGGGCGGGATATGTACGTGGACCAGGAACTGGACATCAACCGGCTGTCCGACTAC
GATGTGGACCATATCGTGCCTCAGAGCTTTCTGAAGGACGACTCCATCGACAACAAGGTGCTGACCAGAAGCGACAA
GAACCGGGGCAAGAGCGACAACGTGCCCTCCGAAGAGGTCGTGAAGAAGATGAAGAACTACTGGCGGCAGCTGCTGA
ACGCCAAGCTGATTACCCAGAGAAAGTTCGACAATCTGACCAAGGCCGAGAGAGGCGGCCTGAGCGAACTGGATAAG
GCCGGCTTCATCAAGAGACAGCTGGTGGAAACCCGGCAGATCACAAAGCACGTGGCACAGATCCTGGACTCCCGGAT
GAACACTAAGTACGACGAGAATGACAAGCTGATCCGGGAAGTGAAAGTGATCACCCTGAAGTCCAAGCTGGTGTCCG
ATTTCCGGAAGGATTTCCAGTTTTACAAAGTGCGCGAGATCAACAACTACCACCACGCCCACGACGCCTACCTGAAC
GCCGTCGTGGGAACCGCCCTGATCAAAAAGTACCCTAAGCTGGAAAGCGAGTTCGTGTACGGCGACTACAAGGTGTA
CGACGTGCGGAAGATGATCGCCAAGAGCGAGCAGGAAATCGGCAAGGCTACCGCCAAGTACTTCTTCTACAGCAACA
TCATGAACTTTTTCAAGACCGAGATTACCCTGGCCAACGGCGAGATCCGGAAGCGGCCTCTGATCGAGACAAACGGC
GAAACCGGGGAGATCGTGTGGGATAAGGGCCGGGATTTTGCCACCGTGCGGAAAGTGCTGAGCATGCCCCAAGTGAA
TATCGTGAAAAAGACCGAGGTGCAGACAGGCGGCTTCAGCAAAGAGTCTATCCTGCCCAAGAGGAACAGCGATAAGC
TGATCGCCAGAAAGAAGGACTGGGACCCTAAGAAGTACGGCGGCTTCGACAGCCCCACCGTGGCCTATTCTGTGCTG
GTGGTGGCCAAAGTGGAAAAGGGCAAGTCCAAGAAACTGAAGAGTGTGAAAGAGCTGCTGGGGATCACCATCATGGA
AAGAAGCAGCTTCGAGAAGAATCCCATCGACTTTCTGGAAGCCAAGGGCTACAAAGAAGTGAAAAAGGACCTGATCA
TCAAGCTGCCTAAGTACTCCCTGTTCGAGCTGGAAAACGGCCGGAAGAGAATGCTGGCCTCTGCCGGCGAACTGCAG
AAGGGAAACGAACTGGCCCTGCCCTCCAAATATGTGAACTTCCTGTACCTGGCCAGCCACTATGAGAAGCTGAAGGG
CTCCCCCGAGGATAATGAGCAGAAACAGCTGTTTGTGGAACAGCACAAGCACTACCTGGACGAGATCATCGAGCAGA
TCAGCGAGTTCTCCAAGAGAGTGATCCTGGCCGACGCTAATCTGGACAAAGTGCTGTCCGCCTACAACAAGCACCGG
GATAAGCCCATCAGAGAGCAGGCCGAGAATATCATCCACCTGTTTACCCTGACCAATCTGGGAGCCCCTGCCGCCTT
CAAGTACTTTGACACCACCATCGACCGGAAGAGGTACACCAGCACCAAAGAGGTGCTGGACGCCACCCTGATCCACC
AGAGCATCACCGGCCTGTACGAGACACGGATCGACCTGTCTCAGCTGGGAGGCGACAAGCGTCCTGCTGCTACTAAG
AAAGCTGGTCAAGCTAAGAAAAAGAAAGCTAGCGGCAGCGGCGCCACCAACTTCAGCCTGCTGAAGCAGGCCGGCGA
CGTGGAGGAGAACCCCGGCCCC**ATG**GCCAAGCCTTTGTCTCAAGAAGAATCCACCCTCATTGAAAGAGCAACGGCTA
CAATCAACAGCATCCCCATCTCTGAAGACTACAGCGTCGCCAGCGCAGCTCTCTCTAGCGACGGCCGCATCTTCACT
GGTGTCAATGTATATCATTTTACTGGGGGACCTTGTGCAGAACTCGTGGTGCTGGGCACTGCTGCTGCTGCGGCAGC
TGGCAACCTGACTTGTATCGTCGCGATCGGAAATGAGAACAGGGGCATCTTGAGCCCCTGCGGACGGTGCCGACAGG
TGCTTCTCGATCTGCATCCTGGGATCAAAGCCATAGTGAAGGACAGTGATGGACAGCCGACGGCAGTTGGGATTCGT
GAATTGCTGCCCTCTGGTTATGTGTGGGAGGGC**TAA**ACGCGTTAAGTCGACAATCAACCTCTGGATTACAAAATTTG
TGAAAGATTGACTGGTATTCTTAACTATGTTGCTCCTTTTACGCTATGTGGATACGCTGCTTTAATGCCTTTGTATC
ATGCTATTGCTTCCCGTATGGCTTTCATTTTCTCCTCCTTGTATAAATCCTGGTTGCTGTCTCTTTATGAGGAGTTG

TGGCCCGTTGTCAGGCAACGTGGCGTGGTGTGCACTGTGTTTGCTGACGCAACCCCCACTGGTTGGGGCATTGCCAC
CACCTGTCAGCTCCTTTCCGGGACTTTCGCTTTCCCCCTCCCTATTGCCACGGCGGAACTCATCGCCGCCTGCCTTG
CCCGCTGCTGGACAGGGGCTCGGCTGTTGGGCACTGACAATTCCGTGGTGTTGTCGGGGAAATCATCGTCCTTTCCT
TGGCTGCTCGCCTGTGTTGCCACCTGGATTCTGCGCGGGACGTCCTTCTGCTACGTCCCTTCGGCCCTCAATCCAGC
GGACCTTCCTTCCCGCGGCCTGCTGCCGGCTCTGCGGCCTCTTCCGCGTCTTCGCCTTCGCCCTCAGACGAGTCGGA
TCTCCCTTTGGGCCGCCTCCCCGCGTCGACTTTAAGACCAATGACTTACAAGGCAGCTGTAGATCTTAGCCACTTTT
TAAAAGAAAAGGGGGGACTGGAAGGGCTAATTCACTCCCAACGAAGACAAGATCTGCTTTTTGCTTGTACTGGGTCT
CTCTGGTTAGACCAGATCTGAGCCTGGGAGCTCTCTGGCTAACTAGGGAACCCACTGCTTAAGCCTCAATAAAGCTT
GCCTTGAGTGCTTCAAGTAGTGTGTGCCCGTCTGTTGTGTGACTCTGGTAACTAGAGATCCCTCAGACCCTTTTAGT
CAGTGTGGAAAATCTCTAGCAGTACGTATAGTAGTTCATGTCATCTTATTATTCAGTATTTATAACTTGCAAAGAAA
TGAATATCAGAGAGTGAGAGGAACTTGTTTATTGCAGCTTATAATGGTTACAAATAAAGCAATAGCATCACAAATTT
CACAAATAAAGCATTTTTTTCACTGCATTCTAGTTGTGGTTTGTCCAAACTCATCAATGTATCTTATCATGTCTGGC
TCTAGCTATCCCGCCCCTAACTCCGCCCATCCCGCCCCTAACTCCGCCCAGTTCCGCCCATTCTCCGCCCCATGGCT
GACTAATTTTTTTTATTTATGCAGAGGCCGAGGCCGCCTCGGCCTCTGAGCTATTCCAGAAGTAGTGAGGAGGCTTT
TTTGGAGGCCTAGGGACGTACCCAATTCGCCCTATAGTGAGTCGTATTACGCGCGCTCACTGGCCGTCGTTTTACAA
CGTCGTGACTGGGAAAACCCTGGCGTTACCCAACTTAATCGCCTTGCAGCACATCCCCCTTTCGCCAGCTGGCGTAA
TAGCGAAGAGGCCCGCACCGATCGCCCTTCCCAACAGTTGCGCAGCCTGAATGGCGAATGGGACGCGCCCTGTAGCG
GCGCATTAAGCGCGGCGGGTGTGGTGGTTACGCGCAGCGTGACCGCTACACTTGCCAGCGCCCTAGCGCCCGCTCCT
TTCGCTTTCTTCCCTTCCTTTCTCGCCACGTTCGCCGGCTTTCCCCGTCAAGCTCTAAATCGGGGGCTCCCTTTAGG
GTTCCGATTTAGTGCTTTACGGCACCTCGACCCCAAAAAACTTGATTAGGGTGATGGTTCACGTAGTGGGCCATCGC
CCTGATAGACGGTTTTTCGCCCTTTGACGTTGGAGTCCACGTTCTTTAATAGTGGACTCTTGTTCCAAACTGGAACA
ACACTCAACCCTATCTCGGTCTATTCTTTTGATTTATAAGGGATTTTGCCGATTTCGGCCTATTGGTTAAAAAATGA
GCTGATTTAACAAAAATTTAACGCGAATTTTAACAAAATATTAACGCTTACAATTTAGGTGGCACTTTTCGGGGAAA
TGTGCGCGGAACCCCTATTTGTTTATTTTTCTAAATACATTCAAATATGTATCCGCTCATGAGACAATAACCCTGAT
AAATGCTTCAATAATATTGAAAAAGGAAGAGTATGAGTATTCAACATTTCCGTGTCGCCCTTATTCCCTTTTTTGCG
GCATTTTGCCTTCCTGTTTTTGCTCACCCAGAAACGCTGGTGAAAGTAAAAGATGCTGAAGATCAGTTGGGTGCACG
AGTGGGTTACATCGAACTGGATCTCAACAGCGGTAAGATCCTTGAGAGTTTTCGCCCCGAAGAACGTTTTCCAATGA
TGAGCACTTTTAAAGTTCTGCTATGTGGCGCGGTATTATCCCGTATTGACGCCGGGCAAGAGCAACTCGGTCGCCGC
ATACACTATTCTCAGAATGACTTGGTTGAGTACTCACCAGTCACAGAAAAGCATCTTACGGATGGCATGACAGTAAG
AGAATTATGCAGTGCTGCCATAACCATGAGTGATAACACTGCGGCCAACTTACTTCTGACAACGATCGGAGGACCGA
AGGAGCTAACCGCTTTTTTGCACAACATGGGGGATCATGTAACTCGCCTTGATCGTTGGGAACCGGAGCTGAATGAA
GCCATACCAAACGACGAGCGTGACACCACGATGCCTGTAGCAATGGCAACAACGTTGCGCAAACTATTAACTGGCGA
ACTACTTACTCTAGCTTCCCGGCAACAATTAATAGACTGGATGGAGGCGGATAAAGTTGCAGGACCACTTCTGCGCT
CGGCCCTTCCGGCTGGCTGGTTTATTGCTGATAAATCTGGAGCCGGTGAGCGTGGGTCTCGCGGTATCATTGCAGCA
CTGGGGCCAGATGGTAAGCCCTCCCGTATCGTAGTTATCTACACGACGGGGAGTCAGGCAACTATGGATGAACGAAA
TAGACAGATCGCTGAGATAGGTGCCTCACTGATTAAGCATTGGTAACTGTCAGACCAAGTTTACTCATATATACTTT
AGATTGATTTAAAACTTCATTTTTAATTTAAAAGGATCTAGGTGAAGATCCTTTTTGATAATCTCATGACCAAAATC
CCTTAACGTGAGTTTTCGTTCCACTGAGCGTCAGACCCCGTAGAAAAGATCAAAGGATCTTCTTGAGATCCTTTTTT
TCTGCGCGTAATCTGCTGCTTGCAAACAAAAAAACCACCGCTACCAGCGGTGGTTTGTTTGCCGGATCAAGAGCTAC
CAACTCTTTTTCCGAAGGTAACTGGCTTCAGCAGAGCGCAGATACCAAATACTGTTCTTCTAGTGTAGCCGTAGTTA
GGCCACCACTTCAAGAACTCTGTAGCACCGCCTACATACCTCGCTCTGCTAATCCTGTTACCAGTGGCTGCTGCCAG
TGGCGATAAGTCGTGTCTTACCGGGTTGGACTCAAGACGATAGTTACCGGATAAGGCGCAGCGGTCGGGCTGAACGG
GGGGTTCGTGCACACAGCCCAGCTTGGAGCGAACGACCTACACCGAACTGAGATACCTACAGCGTGAGCTATGAGAA
AGCGCCACGCTTCCCGAAGGGAGAAAGGCGGACAGGTATCCGGTAAGCGGCAGGGTCGGAACAGGAGAGCGCACGAG
GGAGCTTCCAGGGGGAAACGCCTGGTATCTTTATAGTCCTGTCGGGTTTCGCCACCTCTGACTTGAGCGTCGATTTT
TGTGATGCTCGTCAGGGGGGCGGAGCCTATGGAAAAACGCCAGCAACGCGGCCTTTTTACGGTTCCTGGCCTTTTGC
TGGCCTTTTGCTCACATGTTCTTTCCTGCGTTATCCCCTGATTCTGTGGATAACCGTATTACCGCCTTTGAGTGAGC
TGATACCGCTCGCCGCAGCCGAACGACCGAGCGCAGCGAGTCAGTGAGCGAGGAAGCGGAAGAGCGCCCAATACGCA
AACCGCCTCTCCCCGCGCGTTGGCCGATTCATTAATGCAGCTGGCACGACAGGTTTCCCGACTGGAAAGCGGGCAGT

GAGCGCAACGCAATTAATGTGAGTTAGCTCACTCATTAGGCACCCCAGGCTTTACACTTTATGCTTCCGGCTCGTAT
GTTGTGTGGAATTGTGAGCGGATAACAATTTCACACAGGAAACAGCTATGACCATGATTACGCCAAGCGCGCAATTA
ACCCTCACTAAAGGGAACAAAAGCTGGAGCTGCAAGC
